# *Lactococcus lactis* subsp. cremoris Reprograms the Gut-Liver Axis and Protects Against Liver Injury

**DOI:** 10.64898/2026.09.16.752188

**Authors:** C. Anthony Gacasan, Jaclyn Weinberg, Gabrielle M. Webster, Crystal R. Naudin, ViLinh Tran, Lauren C. Askew, Maria E. Barbian, Dean P. Jones, Rheinallt M. Jones

**Author notes:** **Corresponding author:** Rheinallt M. Jones, Division of Gastroenterology, Hepatology, and Nutrition, Department of Pediatrics, Emory University School of Medicine, 615 Michael Street, Atlanta GA, 30322.

## Abstract

Metabolic diseases are increasingly linked to dysregulation of the gut-liver axis, highlighting the therapeutic potential of probiotics. *Lactococcus lactis* subsp. cremoris (LLC) protects against experimental hepatic steatosis, but the mechanisms underlying its benefits remain poorly understood. Here, we integrated untargeted metabolomics, gnotobiotic mouse models, targeted bile acid profiling, and liver injury paradigms to systematically define LLC-mediated metabolic reprogramming. LLC extensively remodeled serum and hepatic metabolomes in Western diet-fed mice, enriching pathways associated with lipid metabolism, xenobiotic biotransformation, and redox homeostasis. LLC attenuated ethanol-induced steatosis and acetaminophen-induced liver injury, accompanied by activation of Nrf2-dependent antioxidant programs and FXR signaling. LLC monocolonization was sufficient to reprogram the hepatic metabolome, revealing direct host metabolic effects. Reduced intestinal bile acid conjugation emerged as a prominent LLC-associated phenotype. Together, these findings identify LLC as a probiotic that reprograms the gut-liver metabolic axis to enhance metabolic resilience and hepatoprotection.

## Introduction

The gut microbiome exerts profound effects on host metabolism, extending beyond the intestine to shape systemic physiology ^1–4^.

Among extraintestinal organs, the liver occupies a unique position at the interface of dietary, drug-derived, and microbial signals, receiving portal blood enriched in microbial metabolites, bile acids, and xenobiotics ^1–3, 5, 6^. As a result, hepatic metabolic and stress-response pathways are highly sensitive to microbial inputs. Commensal bacteria can influence hepatic antioxidant tone through activation of nuclear factor erythroid 2–related factor 2 (NRF2), a master regulator of cytoprotective, antioxidant, and xenobiotic-response programs ^7–10^. In particular, gut-resident lactobacilli have been shown to activate hepatic NRF2 signaling and protect against oxidative liver injury through microbially derived small molecules ^11^.

Building upon this paradigm, our laboratory identified *Lactococcus lactis* subsp. *cremoris* (LLC) as a potent cytoprotective beneficial microbe ^12^. LLC robustly activated intestinal epithelial NRF2 signaling and conferred resistance to oxidative injury, establishing NRF2-associated intestinal cytoprotection as a defining feature of this strain ^12^. Subsequent studies demonstrated that LLC exerts systemic and context-dependent metabolic effects in vivo, including attenuation of Western-style diet–induced weight gain in female mice without affecting weight gain under control diet conditions ^13^, as well as broader remodeling of systemic metabolism in association with protection from myocardial injury ^14^. Collectively, these findings suggest that the biological effects of LLC extend beyond the intestinal epithelium and may influence metabolic responses in distant organs.

Although NRF2 provides one potential link between microbial activity and host cytoprotection, the liver integrates multiple microbiome-sensitive signaling pathways that govern metabolic homeostasis. Among these are bile acid signaling networks and nuclear receptors including the farnesoid X receptor (FXR) and aryl hydrocarbon receptor (AHR) ^15–18^. Beyond their classical role in lipid absorption, bile acids function as signaling molecules that regulate hepatic lipid and glucose metabolism, xenobiotic processing, inflammatory tone, and oxidative stress responses ^17, 19, 20^. FXR, whose activity is influenced by the composition of the bile acid pool, coordinates bile acid synthesis, transport, and metabolic homeostasis ^15–17, 21^. Microbial metabolism can substantially reshape this signaling environment through bile acid deconjugation and subsequent transformations, thereby altering the intestinal bile acid pool and its interaction with host receptors ^19, 20, 22^. In parallel, microbiome-derived tryptophan metabolites can engage AHR and influence host transcriptional programs ^18, 23^. Consistent with these relationships, germ-free and conventionally colonized animals exhibit substantial differences in both bile acid metabolism and systemic metabolite profiles, underscoring the sensitivity of hepatic and enterohepatic metabolism to microbial composition ^4, 22, 24^.

Here, we sought to determine whether LLC, previously characterized by intestinal cytoprotective activity and systemic metabolic effects, is associated with alterations in hepatic metabolism and susceptibility to liver injury ^12–14^. Using untargeted metabolomics across conventional and gnotobiotic mouse models, we examined the effects of LLC on hepatic metabolic organization and identified changes involving xenobiotic, amino acid, lipid, and mitochondrial-associated pathways. We further tested the functional relevance of these metabolic changes using acute acetaminophen- and ethanol-induced liver injury models and evaluated hepatic transcriptional programs related to antioxidant defense and xenobiotic metabolism.

Comparative analyses in gnotobiotic and conventionally colonized mice were then used to identify LLC-associated metabolic features that were reproducible across distinct microbial contexts. Network-based analysis further highlighted coordinated metabolic programs involving xenobiotic metabolism, mitochondrial function, lipid oxidation, and bile acid metabolism, providing a rationale for targeted examination of bile acid composition. Because bile acid deconjugation represents a major microbial transformation within the enterohepatic system ^25–27^, we specifically investigated whether LLC could alter bile acid conjugation both in isolation and within a complex intestinal microbiota.

Targeted bile acid profiling revealed that LLC monocolonization was sufficient to reduce fecal bile acid conjugation, including reciprocal changes in conjugated and unconjugated primary bile acids. A similar reduction in conjugated fecal bile acids was observed following LLC administration to conventionally colonized mice, demonstrating that this phenotype persists in the presence of a complex microbiota. In contrast, corresponding class-level hepatic bile acid pools were comparatively stable, indicating that LLC-associated bile acid remodeling is substantially more pronounced within the intestinal compartment. These changes occurred alongside alterations in hepatic NRF2-associated antioxidant genes, *Fxr*, and *Cyp2e1*, although the present experiments do not establish a causal relationship between intestinal bile acid remodeling and hepatic transcriptional or injury phenotypes. To examine whether probiotic-derived factors could engage related transcriptional responses in a human hepatic cell model, HepG2 cells were exposed to bacterial culture supernatants from LLC or LGG. Supernatants from both strains significantly increased *GCLC* expression, whereas *CYP2E1* expression was not significantly altered, indicating that soluble probiotic-derived factors can induce antioxidant-associated transcriptional responses in a human hepatocyte-derived cell model. Together, these findings expand the biological scope of LLC beyond intestinal cytoprotection and identify altered intestinal bile acid conjugation as one component of a broader LLC-associated metabolic program linking the gut microbiota with hepatic metabolism, stress-response pathways, and susceptibility to injury.

## Results

### LLC Remodels the Hepatic and Circulating Metabolome in Western Diet–Fed Mice

Given that LLC exhibits pronounced metabolic effects under Western-style diet conditions, we performed untargeted LC–MS metabolomics on serum and liver from Western diet–fed mice supplemented with vehicle (HBSS), LGG, or LLC (n = 10 per group) to define metabolic pathways associated with LLC treatment ^13^. Multivariate analysis demonstrated substantial treatment-associated metabolic remodeling. Partial least squares–discriminant analysis (PLS-DA) showed separation between LLC- and HBSS-treated mice in both serum (**Figure 1B**) and liver (**Figure 1C**). Consistent with these patterns, PERMANOVA demonstrated significant differences between LLC and HBSS groups in serum (R² = 0.363, *P* = 0.001, adjusted *P* = 0.003), whereas LGG- and LLC-treated mice did not significantly differ (R² = 0.131, *P* = 0.104; **Supplementary Table 1A**). In liver, LLC-treated mice similarly differed from HBSS controls (R² = 0.356, *P* = 0.001, adjusted *P* = 0.003), with significant separation also observed between LLC and LGG groups (R² = 0.202, *P* = 0.009, adjusted *P* = 0.0135; **Supplementary Table 1B**). Thus, treatment group accounted for approximately 35–36% of the metabolomic variation observed in pairwise LLC-versus-HBSS comparisons in both serum and liver, supporting a substantial systemic metabolic response to LLC under Western diet conditions.

**Figure 1.**
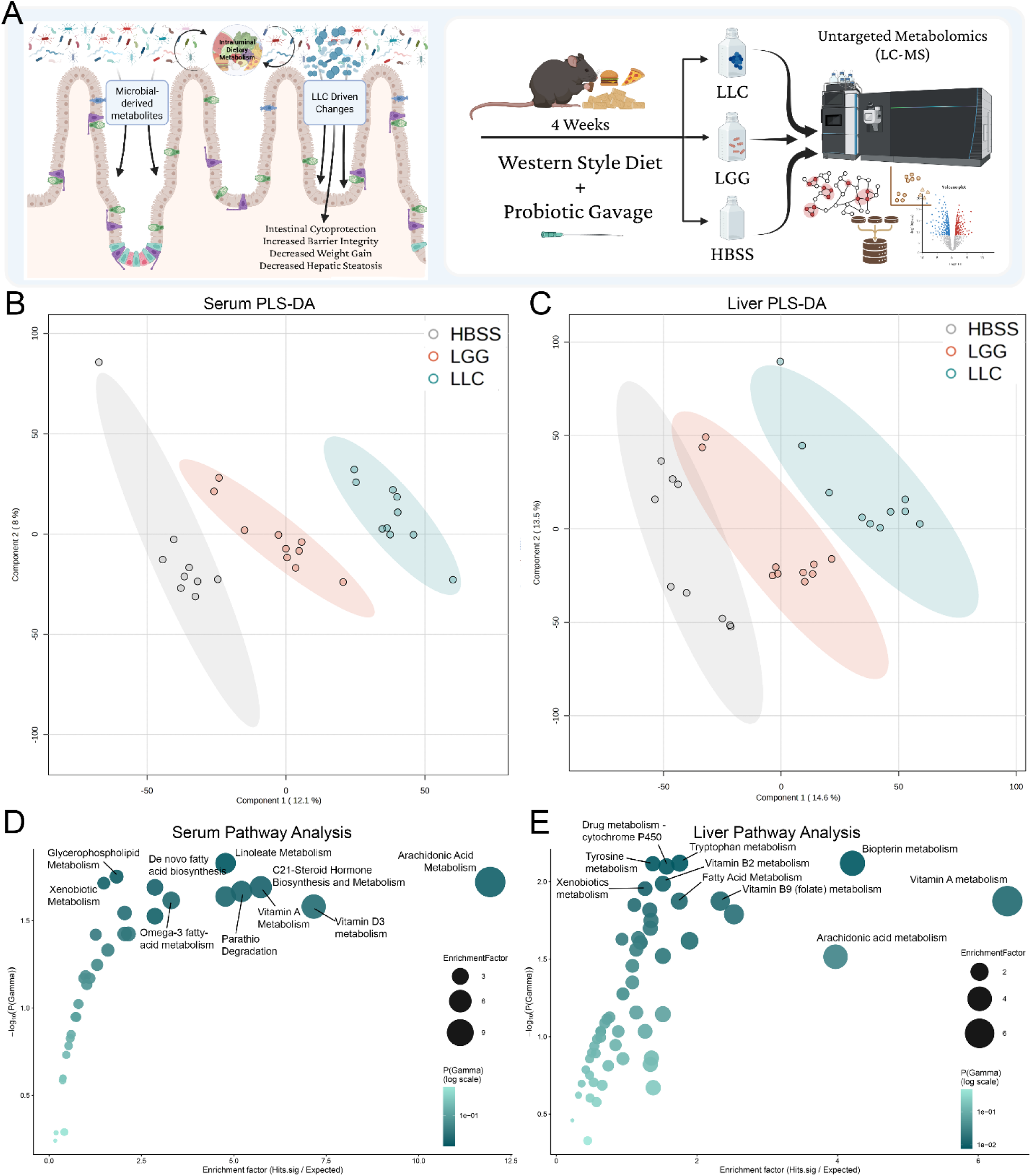
Serum and Liver Metabolomic Profiling in Western Diet–Fed Mice Treated with LLC (n = 10 per group). **(A)** Proposed model illustrating how LLC-driven changes in intestinal, microbiota-derived metabolites promote cytoprotective, metabolic, and hepatoprotective effects (left), and experimental schema depicting four weeks of probiotic administration during Western-style diet feeding followed by untargeted serum and liver metabolomic analyses (right). **(B-C)** Partial least squares– discriminant analysis (PLS-DA) of untargeted metabolomics from serum (B) and liver (C) of Western diet–fed mice treated with HBSS, LGG, or LLC. Each point represents an individual mouse (n = 10 per group). **(D–E)** Pathway enrichment analysis performed using mummichog from serum (C) and liver (D) metabolomic datasets, displayed as bubble plots. Bubble size represents enrichment factor (Hits.sig/Expected), and the y-axis indicates −log10(p-value).

To identify metabolic pathways associated with these differences, we performed pathway enrichment analysis using mummichog ^28^. In serum, the most significantly enriched pathway in LLC-treated mice was linoleate metabolism (enrichment factor = 4.78, *P* = 7.04 × 10⁻⁴), followed by glycerophospholipid metabolism (enrichment factor = 1.83, *P* = 0.0177) and arachidonic acid metabolism (enrichment factor = 11.94, *P* = 0.0190), indicating broad alterations in lipid-associated metabolic pathways (**Figure 1D**). Xenobiotic metabolism was also enriched in LLC-treated mice (enrichment factor = 1.48, *P* = 0.0193), suggesting additional effects on circulating metabolites associated with xenobiotic processing.

In liver, pathway enrichment revealed a more focused signature involving amino acid and xenobiotic metabolism (**Figure 1E**). Tryptophan metabolism was the most significantly enriched pathway (enrichment factor = 1.76, *P* = 7.6 × 10⁻⁵), followed by biopterin metabolism (enrichment factor = 4.21, *P* = 6.0 × 10⁻⁴) and tyrosine metabolism (enrichment factor = 1.37, *P* = 0.0031). Drug metabolism–cytochrome P450 was also significantly enriched (enrichment factor = 1.57, *P*= 0.0183), implicating hepatic xenobiotic-associated metabolic pathways ^7, 8^. Collectively, these findings demonstrate that LLC treatment is associated with substantial remodeling of the circulating and hepatic metabolomes under Western diet conditions, including alterations in serum lipid-associated pathways and hepatic amino acid and cytochrome P450–associated metabolism. These metabolic signatures extend previous observations of systemic LLC-associated metabolic remodeling and provide a biological framework for examining whether LLC influences hepatic responses to metabolic and xenobiotic stress ^13, 14^.

### LLC Attenuates Alcohol-Induced Hepatic Microsteatosis

To determine whether LLC modifies the hepatic response to acute alcohol exposure, male mice were pretreated with vehicle (HBSS), LGG, or LLC by daily gavage for 2 weeks (n = 10 per group) prior to acute ethanol challenge (6 g/kg). Hepatic lipid accumulation was assessed by Oil Red O staining with hematoxylin counterstain (**Figure 2A–B**), a measure of hepatic neutral lipid accumulation following acute ethanol exposure ^29^. Compared with calorie-matched dextrose controls, HBSS-treated mice exhibited marked hepatic lipid accumulation following ethanol challenge. Two-way ANOVA of log-transformed Oil Red O quantification demonstrated a significant Exposure × Microbe interaction (*P* = 3.6 × 10⁻⁵), indicating that the effect of ethanol on hepatic lipid accumulation differed according to probiotic treatment.

**Figure 2.**
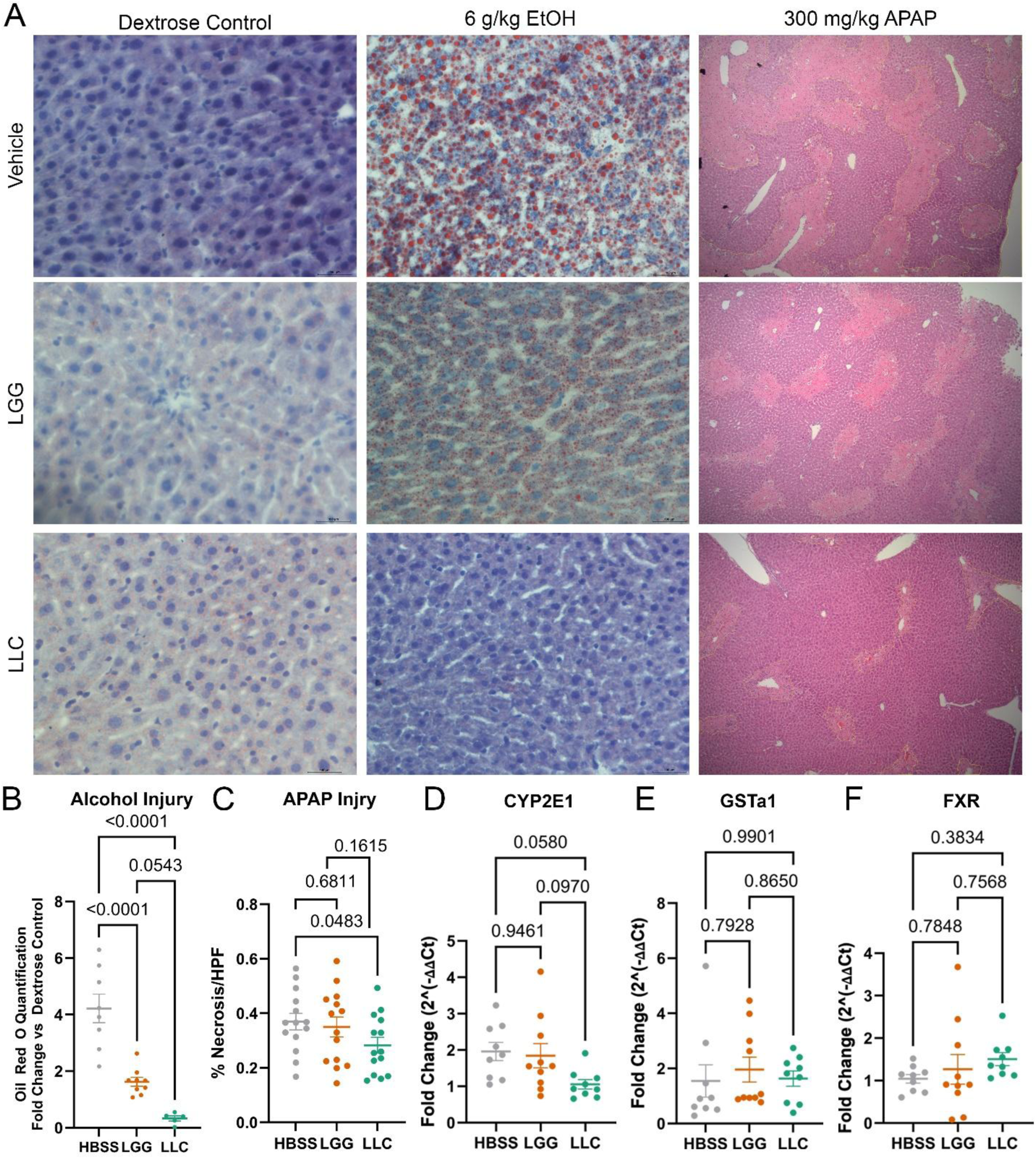
LLC Protects Against Alcohol- and Acetaminophen-Induced Liver Injury (n = 10 per group). All mice were male and pretreated with HBSS vehicle, LGG, or LLC for 2 weeks prior to injury challenge (n = 10 per group). **(A)** Representative histological images of liver sections following injury challenge. The first and second columns correspond to the alcohol injury model (6 g/kg challenge) and include Oil Red O staining with H&E counterstain. The first column represents calorie-matched dextrose control, and the second column represents alcohol challenge. The third column shows acetaminophen (APAP) challenge (300 mg/kg) with H&E staining; centrilobular necrosis is outlined. Rows represent treatment groups: HBSS (top), LGG (middle), and LLC (bottom). Images are representative. **(B)** Quantification of Oil Red O staining from the alcohol injury model. **(C)** Quantification of percent centrilobular necrosis per high-powered field in APAP-challenged livers. **(D–F)** Hepatic mRNA expression of **(D)** CYP2E1, **(E)** GSTa1, and **(F)** FXR measured by RT-qPCR in APAP-treated mice.

To further compare the magnitude of the ethanol response among treatment groups, ethanol values were normalized to their respective dextrose controls and analyzed by one-way ANOVA. Fold induction differed significantly among groups (F = 30.80, *P* < 0.0001, R² = 0.7643). Tukey’s multiple-comparisons testing demonstrated significantly lower ethanol-induced lipid accumulation in both LGG-treated (mean difference versus HBSS = 2.596, *P* < 0.0001) and LLC-treated mice (mean difference versus HBSS = 3.889, *P* < 0.0001). Although the reduction was numerically greater following LLC treatment than LGG treatment, the difference between the two probiotic groups did not reach statistical significance (*P* = 0.0543). Together, these findings demonstrate that LLC pretreatment attenuates acute ethanol-induced hepatic lipid accumulation, an effect that was also observed following LGG treatment.

### LLC Reduces APAP-Induced Centrilobular Necrosis and Modulates Xenobiotic Metabolic Gene Expression

We next assessed whether LLC modifies susceptibility to acetaminophen (APAP)-induced hepatotoxicity. Male mice were supplemented with vehicle (HBSS), LGG, or LLC by daily gavage for 2 weeks (n = 10 per group) before APAP challenge (300 mg/kg). Liver sections were analyzed by hematoxylin and eosin staining, and centrilobular necrosis was quantified as percent necrotic area per high-powered field (**Figure 2A, C**). HBSS-treated mice exhibited extensive centrilobular necrosis following APAP challenge, whereas LLC-treated mice exhibited significantly less necrotic area than HBSS controls (*P* = 0.04). LGG supplementation did not significantly reduce APAP-induced centrilobular necrosis relative to HBSS.

Because APAP hepatotoxicity is driven in part by cytochrome P450–mediated conversion of APAP to the reactive intermediate *N*-acetyl-*p*-benzoquinone imine (NAPQI), we next examined hepatic expression of genes involved in xenobiotic metabolism and stress-response signaling ^30–32^. *Cyp2e1* transcript abundance was lower in LLC-treated mice relative to HBSS controls, although this difference did not reach statistical significance (*P* = 0.058; **Figure 2D**). *Gsta1* expression showed a similar direction toward increased expression following LLC treatment but was not significantly different from HBSS (**Figure 2E**). Hepatic *Fxr* expression also trended higher in LLC-treated mice without reaching statistical significance (**Figure 2F**).

Together, these data demonstrate that LLC pretreatment reduces APAP-induced centrilobular necrosis and is accompanied by directional changes in hepatic genes associated with xenobiotic metabolism and stress-response signaling. Although these transcriptional changes are consistent with the broader hepatic metabolic signatures identified following LLC treatment, they do not independently establish the mechanism underlying protection from APAP-induced injury.

### Comparative Metabolomic Analysis Identifies LLC-Associated Signatures Across Microbial Contexts

To distinguish hepatic metabolic features associated with LLC from those broadly associated with microbial colonization, we performed comparative metabolomic analysis across conventional and gnotobiotic cohorts maintained on standard mouse chow (**Figure 3A**). Across all seven experimental groups, PERMANOVA demonstrated a significant overall difference in hepatic metabolomic composition (R² = 0.268, F = 2.56, *P* = 0.0001; **Supplementary Table 2A**), with group identity accounting for approximately 27% of variation in the combined metabolomic dataset. Consistent with this overall effect, PCA of all seven groups (**Figure 3B**) demonstrated broad separation among germ-free, monocolonized, SPF-colonized, and conventionally colonized cohorts. We next performed within-context comparisons to define the metabolic effects associated with LLC treatment under conventional and gnotobiotic conditions.

**Figure 3.**
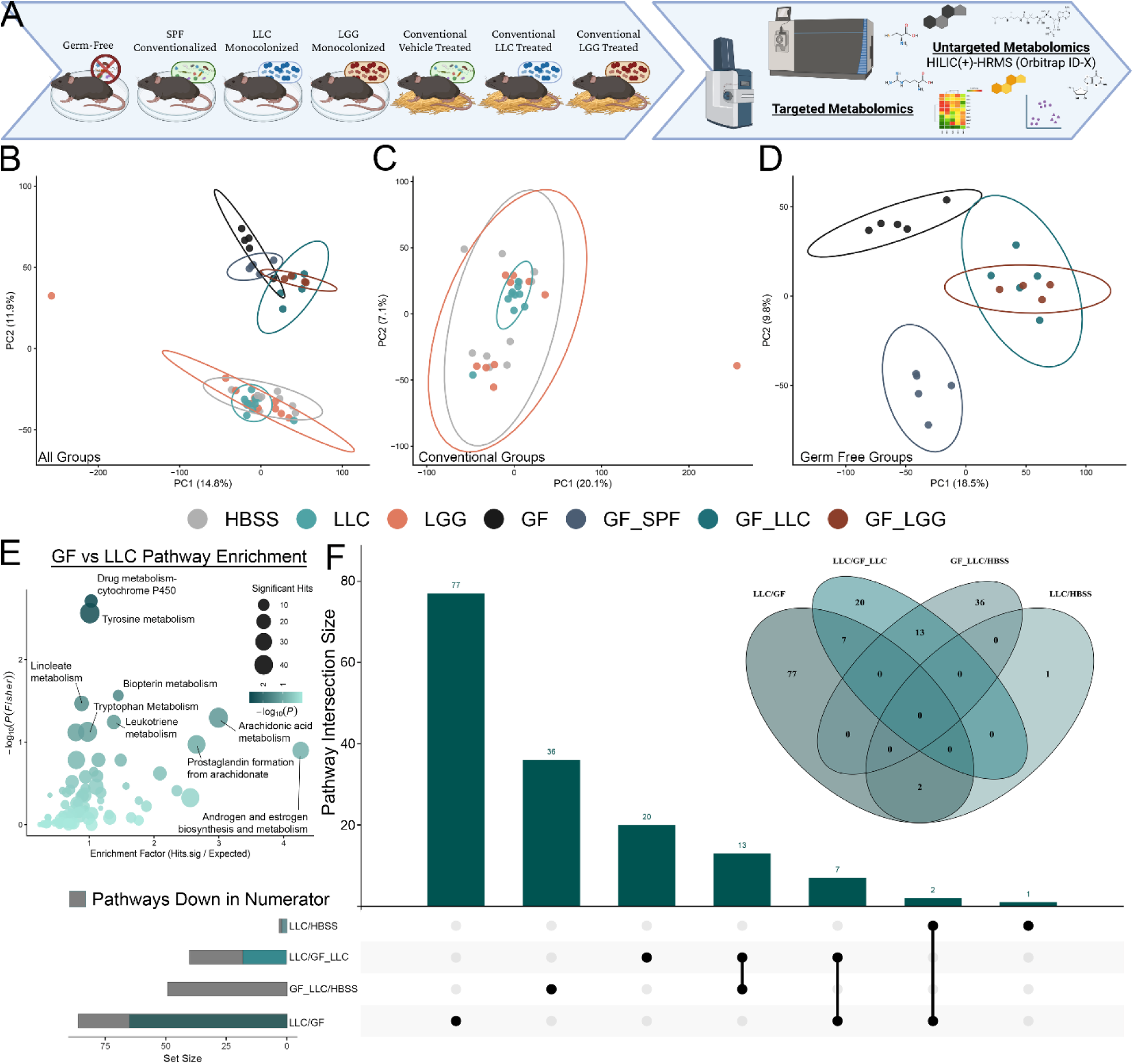
Comparative Metabolomic Analysis Across Conventional and Gnotobiotic Cohorts. Untargeted metabolomics was performed across seven groups: conventional HBSS, LGG, and LLC mice gavaged daily for 4 weeks (n = 10 per group), and germ-free (GF) mice (n = 5 per group) that were either maintained germ-free, colonized with SPF microbiota via fecal gavage and bedding exchange (GF_SPF), monocolonized with LLC by daily gavage for 2 weeks (GF_LLC), or monocolonized with LGG by daily gavage for 2 weeks (GF_LGG). **(A)** Schematic of experimental design **(B)** Principal component analysis (PCA) of all seven groups. **(C)** PCA of conventional mice treated with HBSS, LGG, or LLC (n = 10 per group). **(D)** PCA of germ-free and colonized germ-free groups (GF, GF_SPF, GF_LLC; n = 5 per group, GF=LGG; n=4 per group). **(E)** Pathway enrichment analysis comparing GF and LLC groups. **(F)** Direction-aware UpSet plot summarizing overlap of enriched metabolic pathways across pairwise contrasts (LLC/HBSS, LLC/GF_LLC, GF_LLC/HBSS, LLC/GF), indicating shared and unique features among comparisons. Corresponding Venn diagram above UpSet plot showing overlapping pathways common among indicated comparisons.

Within conventionally colonized mice (HBSS, LGG, and LLC; **Figure 3C**), pairwise PERMANOVA demonstrated significant separation between HBSS- and LLC-treated mice (R² = 0.084, *P* = 0.0012, FDR-adjusted *P* = 0.0029), whereas LLC and LGG did not significantly differ (R² = 0.063, *P* = 0.1125). Similarly, HBSS and LGG did not significantly differ (R² = 0.060, *P* = 0.1830; **Supplementary Table 2B**). These findings indicate that LLC treatment is associated with a modest but statistically significant shift in hepatic metabolomic composition relative to vehicle in conventionally colonized mice.

Within the gnotobiotic cohort, microbial colonization produced more pronounced alterations in hepatic metabolomic composition (**Figure 3D**). GF and GF_LLC mice differed significantly (R² = 0.263, *P* = 0.0081, FDR-adjusted *P* = 0.0106), as did GF and GF_LGG mice (R² = 0.282, *P* = 0.0090, FDR-adjusted *P* = 0.0111). Colonization with an SPF microbiota also significantly altered the metabolome relative to GF controls (GF versus GF_SPF: R² = 0.217, *P* = 0.0076). GF_LLC and GF_LGG mice did not significantly differ from one another (R² = 0.125, *P* = 0.4607; **Supplementary Table 2B**), indicating substantial overlap in the hepatic metabolic responses elicited by the two monocolonizing strains. Together, these findings demonstrate that colonization with LLC alone is sufficient to alter hepatic metabolomic organization in the absence of a complex microbial community.

Pathway enrichment analysis of the GF_LLC versus GF comparison identified metabolic pathways associated with xenobiotic and amino acid metabolism (**Figure 3E**). Drug metabolism– cytochrome P450 was enriched (enrichment factor = 1.03, *P* = 0.00195), implicating xenobiotic-associated metabolism in the hepatic response to LLC monocolonization. Tyrosine metabolism was also enriched (enrichment factor = 1.01, *P* = 0.00273), along with biopterin metabolism (enrichment factor = 1.45, *P* = 0.0272). Linoleate metabolism also showed enrichment (*P* = 0.0339), whereas arachidonic acid metabolism showed a weaker nominal signal (enrichment factor = 2.99, *P* = 0.0506). These findings demonstrate that LLC monocolonization is sufficient to remodel hepatic metabolic pathways, including cytochrome P450–associated xenobiotic metabolism, without requiring the presence of a complex intestinal microbiota.

To determine whether LLC-associated metabolic programs were reproducible across microbial contexts, we next compared directionally annotated pathway sets generated from the relevant pairwise contrasts. Direction-aware pathway overlap was summarized using an UpSet framework (**Figure 3F**) ^33^. Although most enriched pathways were specific to individual experimental contexts, several pathways recurred across comparisons (**Supplementary Tables S3–S5**). We therefore used weighted correlation network analysis to determine whether LLC colonization was associated with coordinated hepatic metabolic programs.

### Network Analysis Identifies LLC-Associated Metabolic Programs

Weighted correlation network analysis (WGCNA) of liver metabolomics from the gnotobiotic cohorts identified coordinated modules of metabolite features associated with microbial colonization state (**Figure 4**) ^34^. Network construction resolved 18 distinct metabolite co-abundance modules, with module–trait correlation analysis revealing distinct patterns across GF, GF_LLC, GF_LGG, and GF_SPF mice. GF mice displayed a pattern of module associations that differed from colonized groups, while monocolonized and SPF-colonized mice exhibited distinct but partially overlapping module–trait relationships. These findings indicate that introduction of individual bacterial strains or a complex microbiota is associated with coordinated reorganization of hepatic metabolic networks.

**Figure 4.**
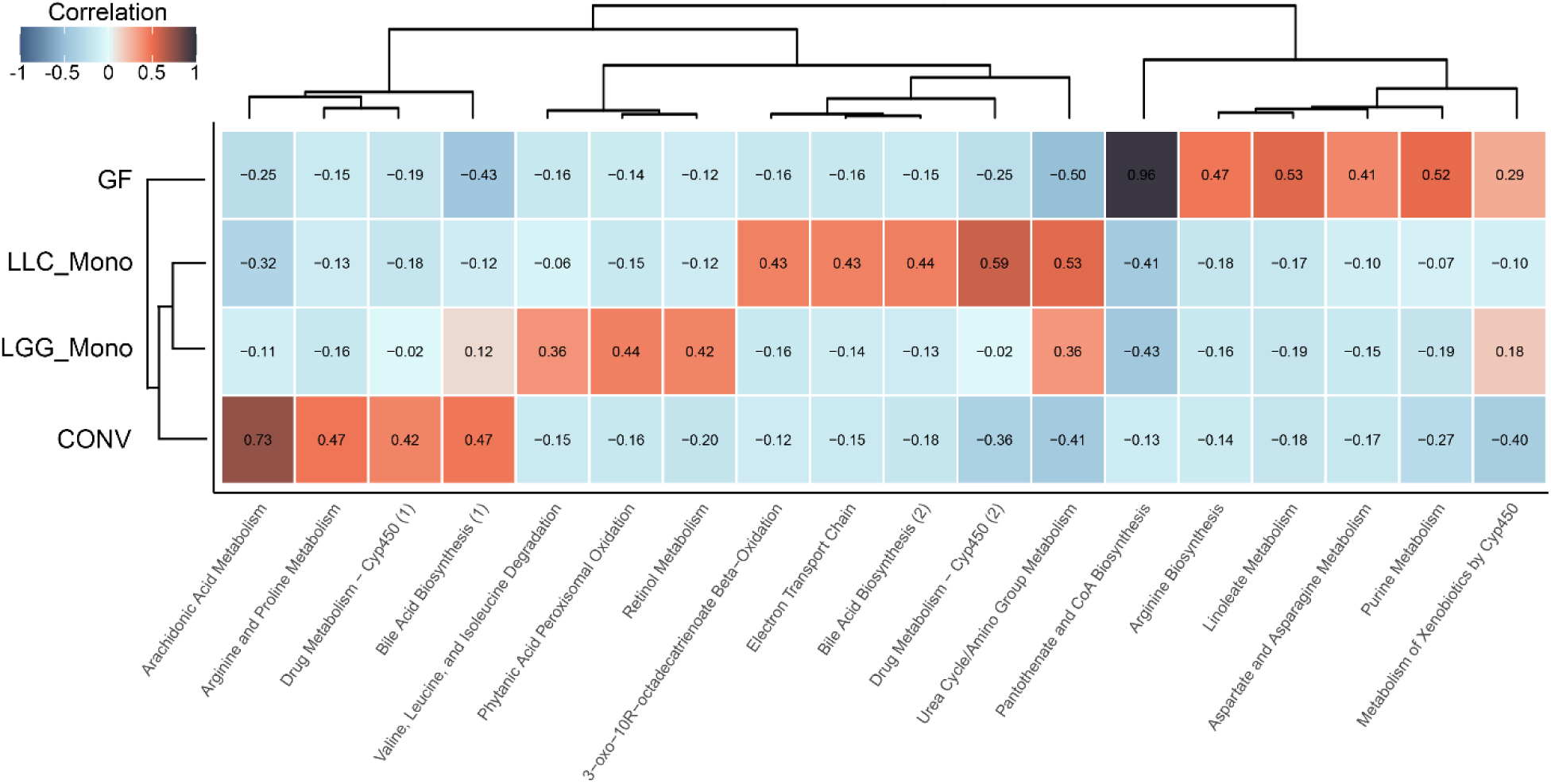
Weighted correlation network analysis (WGCNA) of liver metabolomics identifies LLC-associated metabolic modules in gnotobiotic mice. Hierarchical clustering of liver metabolite features using a minimum module size of 250 identified 18 distinct co-abundance modules across GF mice subjected to different colonization conditions. The module–trait correlation heatmap displays Pearson correlations between module eigengenes and treatment groups (GF, LLC_Mono, LGG_Mono, and conventionalized with SPF microbiota [CONV]), with correlation coefficients shown within each cell. Hierarchical clustering of modules is displayed across the top, and treatment groups are shown along the side. Group sizes were *n* = 5 for GF, LLC_Mono, and CONV and *n* = 4 for LGG_Mono. Metabolite features within each module were subjected to mummichog pathway enrichment analysis, and modules are labeled by their top-ranked enriched pathway, linking module-level co-abundance patterns to putative biological processes. Five modules were significantly associated with LLC monocolonization (*P* < 0.05), including a module significantly enriched for drug metabolism–cytochrome P450, while other LLC-associated modules contained top-ranked annotations related to mitochondrial metabolism, lipid oxidation, and bile acid biosynthesis.

Among the monocolonized groups, five metabolite modules were significantly correlated with the GF_LLC condition (*P* < 0.05), suggesting that LLC colonization is associated with changes across multiple coordinated metabolic programs rather than a single isolated pathway. Hierarchical clustering of module–trait correlation profiles further demonstrated both shared and distinct patterns among GF_LLC, GF_LGG, and GF_SPF mice, consistent with overlapping metabolic effects of microbial colonization together with strain- and community-dependent differences (**Figure 4**).

To investigate the biological processes represented within LLC-associated modules, we performed pathway enrichment analysis using mummichog v2.0 ^28^. Drug metabolism–cytochrome P450 was significantly enriched within one LLC-associated module (*P* = 0.016), providing additional evidence that xenobiotic-associated metabolism is a prominent component of the hepatic response to LLC colonization. Other LLC-associated modules were characterized by top-ranked pathway annotations related to mitochondrial and lipid metabolism, including electron transport chain metabolism (*P* = 0.056) and 3-oxo-10R-octadecatrienoate β-oxidation (*P* = 0.12). Because these latter associations did not meet the predefined threshold for statistical significance, they were considered hypothesis-generating rather than evidence of significant pathway enrichment.

Bile acid biosynthesis was also identified among the top pathway annotations of an LLC-associated module (*P* = 0.14) ^17, 21^. Although this association did not reach statistical significance, the emergence of bile acid metabolism within an LLC-correlated metabolic module, together with established microbiome-dependent regulation of bile acid composition and enterohepatic signaling ^20, 22^, provided a rationale for directly examining bile acid metabolism in subsequent targeted analyses.

### LLC Remodels the Hepatic and Fecal Bile Acid Pool

Because bile acid metabolism emerged among the top pathway annotations within an LLC-associated metabolite module, we next performed targeted bile acid profiling to directly assess whether LLC alters bile acid composition in vivo. To evaluate these effects across distinct microbial contexts, bile acids were measured in seven experimental groups: germ-free mice (GF), germ-free mice monocolonized with LLC (GF_LLC) or LGG (GF_LGG), germ-free mice conventionalized with a complete SPF microbiota (GF_SPF), and conventionally colonized mice treated with vehicle (HBSS), LLC, or LGG. By examining both fecal and hepatic bile acid pools, this design allowed us to determine whether LLC-associated changes in bile acid composition were evident in a reductionist monocolonization model and whether similar changes were retained in the presence of a complex intestinal microbiota (**Figure 5**) ^22, 24^.

**Figure 5.**
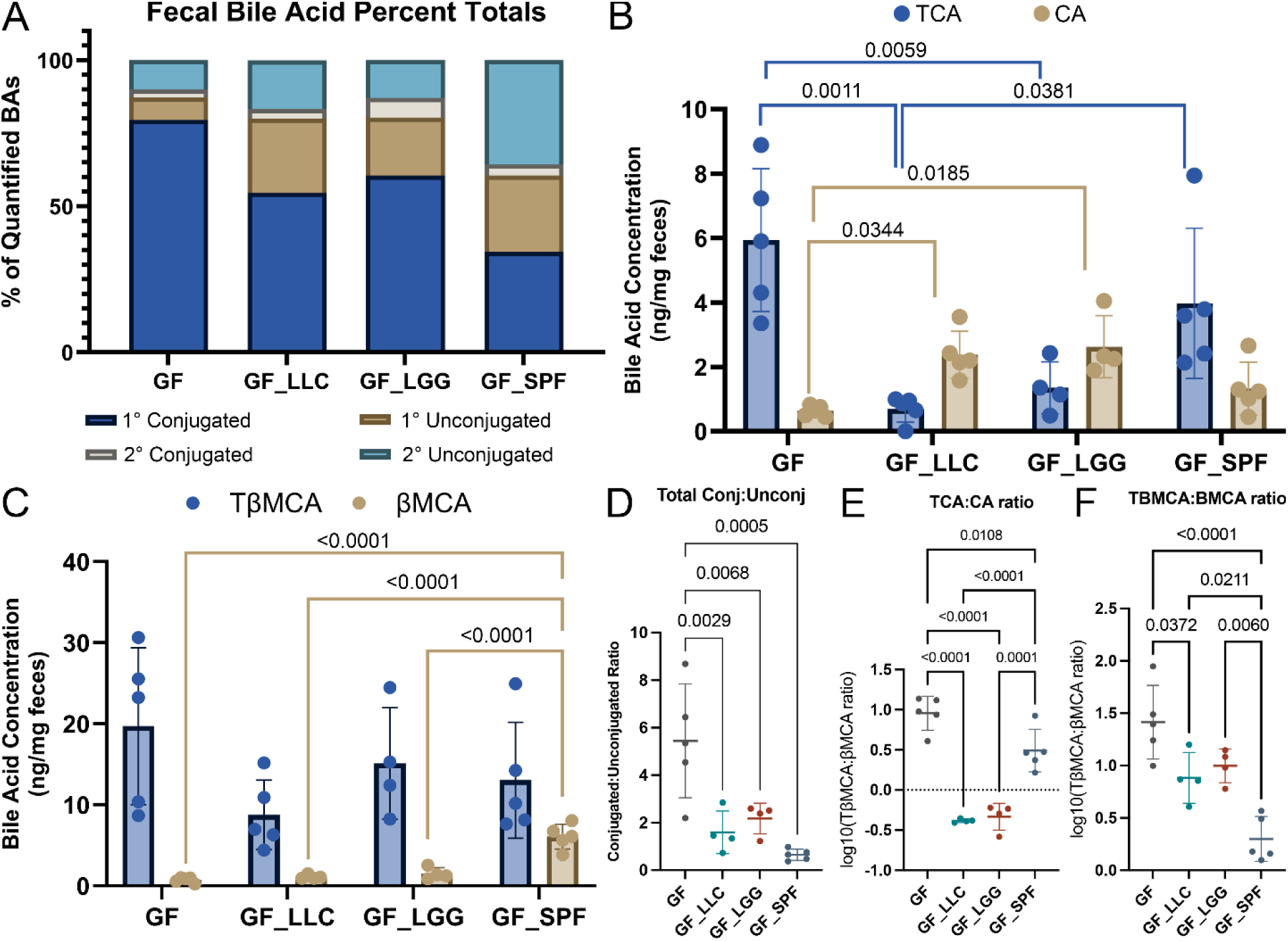
Microbial colonization alters fecal bile acid composition and conjugation in gnotobiotic mice. Fecal bile acid profiles were compared among germ-free mice (GF), mice monocolonized with LLC (GF_LLC) or LGG (GF_LGG), and germ-free mice conventionalized with SPF microbiota (GF_SPF). **(A)** Relative composition of the quantified fecal bile acid pool, shown as the percentage contributed by primary conjugated, secondary conjugated, primary unconjugated, and secondary unconjugated bile acids. **(B)** Fecal concentrations of taurocholic acid (TCA) and its unconjugated counterpart cholic acid (CA). **(C)** Fecal concentrations of tauro-β-muricholic acid (TβMCA) and β-muricholic acid (βMCA). **(D)** Ratio of total conjugated to total unconjugated bile acids. **(E)** Log10-transformed TCA:CA ratio and **(F)** log10-transformed TβMCA:βMCA ratio, illustrating shifts in bile acid conjugation across colonization states. Individual points represent individual mice and summary bars indicate group mean ± SD. GF_LGG with *n* = 5 and all others with *n* = 5 mice per group. Statistical comparisons were performed using one-way ANOVA followed by Tukey’s multiple-comparisons test. Exact Tukey-adjusted p values for the indicated pairwise comparisons are shown above the corresponding brackets.

Targeted bile acid measurements were performed using an ion mobility–enabled LC–MS platform incorporating curated *m/z*, collision cross-section (CCS), and molecular formula information ^35^. The analytical panel included 23 bile acids with authentic standards, permitting absolute quantification when calibration curves were available, while additional bile acid species were detected and assessed semi-quantitatively using matched *m/z* and CCS information. In fecal samples, 70 bile acid species were detected, of which 21 were quantified using authentic standards. In liver, 41 bile acid species were detected, including 18 that were quantified using authentic standards (**Supplementary Tables S6–S7; Supplementary Figures 1–2**). Measurements exceeding the upper limit of the corresponding calibration curve were retained as estimated values for comparative analyses and were flagged accordingly. Subsequent quantitative analyses were restricted to bile acid species with authentic standards and consistent detection across experimental groups.

Because microbial bile salt hydrolase (BSH) activity represents a major route of bile acid deconjugation ^25, 36^, we next assessed whether LLC exhibited bile acid deconjugation capacity in vitro. LLC, LGG, or vehicle control cultures were incubated in MRS medium containing either a mixed pool of conjugated bile acids (0.5% w/v glycocholic acid, glycodeoxycholic acid, taurocholic acid, and taurodeoxycholic acid; **Supplementary Figure 3**) or individual conjugated bile acids supplied at 1 mM (**Supplementary Figure 4**). Visible precipitation was observed in cultures containing LLC and LGG relative to controls, consistent with deconjugation and reduced solubility of the resulting unconjugated bile acids ^25, 36^. Precipitation was qualitatively more pronounced in LLC cultures than in LGG cultures under the conditions tested. These observations demonstrate that LLC possesses bile acid deconjugation capacity in vitro and provided additional rationale for examining bile acid conjugation in vivo. Because this assay was qualitative, these results were not interpreted as a quantitative comparison of BSH activity between LLC and LGG.

### LLC Monocolonization is Sufficient to Remodel Fecal Bile Acid Conjugation

We first used the gnotobiotic cohort to determine whether LLC alone was sufficient to alter intestinal bile acid composition. Fecal bile acids were quantified in GF mice, mice monocolonized with LLC (GF_LLC) or LGG (GF_LGG), and GF mice conventionalized with an SPF microbiota (GF_SPF; **Figure 5**).

GF mice exhibited a fecal bile acid pool dominated by conjugated primary bile acids, consistent with the absence of microbial bile acid deconjugation ^22^. Colonization substantially altered the composition of this pool (**Figure 5A**). GF_SPF mice exhibited the broadest remodeling, characterized by increased representation of unconjugated primary and secondary bile acids and a corresponding reduction in conjugated species. Monocolonization produced a more restricted phenotype, with both GF_LLC and GF_LGG mice exhibiting reduced contributions of conjugated primary bile acids relative to GF controls.

We next examined the major cholic acid conjugated–unconjugated pair (**Figure 5B**). Taurocholic acid (TCA) concentrations were reduced in GF_LLC mice relative to GF controls (*P* = 0.0011) and were similarly reduced in GF_LGG mice (*P* = 0.0059). TCA concentrations were also lower in GF_LLC than in GF_SPF mice (*P* = 0.0381). Conversely, the unconjugated counterpart cholic acid (CA) was increased in both GF_LLC (*P* = 0.0344) and GF_LGG (*P* = 0.0185) relative to GF controls. Thus, monocolonization with either strain produced reciprocal changes in conjugated and unconjugated forms of cholic acid.

A similar pattern was observed for the β-muricholic acid pair (**Figure 5C**). TβMCA concentrations were reduced following monocolonization, whereas βMCA showed a reciprocal increase. GF_SPF mice exhibited a substantially greater increase in βMCA than GF, GF_LLC, or GF_LGG mice (all *P* < 0.0001), consistent with more extensive bile acid remodeling following colonization with a complex microbiota.

To determine whether these species-level changes reflected a broader alteration in bile acid conjugation, we next examined the ratio of total conjugated to unconjugated bile acids (**Figure 5D**). This ratio was significantly reduced in GF_LLC mice relative to GF controls (*P* = 0.0029). GF_LGG mice showed a similar reduction (*P* = 0.0068), while GF_SPF mice exhibited the largest decrease (*P* = 0.0005). These findings indicate that the reciprocal changes observed in individual bile acid species were part of a broader shift toward an unconjugated fecal bile acid pool following microbial colonization.

Consistent with this pattern, the TCA:CA ratio was markedly reduced following LLC monocolonization (*P* < 0.0001 versus GF; **Figure 5E**). A similarly reduced ratio was observed in GF_LGG mice (*P* < 0.0001 versus GF), while GF_SPF mice also differed from GF controls (*P* = 0.0108). The TCA:CA ratio was lower in both GF_LLC and GF_LGG mice than in GF_SPF mice (*P* < 0.0001 and *P* = 0.0001, respectively), demonstrating pronounced remodeling of this conjugated–unconjugated bile acid pair following monocolonization. Finally, analysis of the TβMCA:βMCA ratio demonstrated a directionally concordant response (**Figure 5F**). LLC monocolonization significantly reduced the TβMCA:βMCA ratio relative to GF controls (*P* = 0.0372). GF_SPF mice exhibited an even greater reduction relative to GF (*P* < 0.0001) and also had lower ratios than GF_LLC (*P* = 0.0211) and GF_LGG (*P* = 0.0060), reflecting the pronounced expansion of unconjugated βMCA following conventionalization.

Together, these findings demonstrate that LLC monocolonization is sufficient to reduce fecal bile acid conjugation in vivo, producing reciprocal changes in conjugated and unconjugated primary bile acids and a broader reduction in the conjugated-to-unconjugated bile acid ratio. LGG produced a partially overlapping phenotype, while SPF conventionalization resulted in broader bile acid remodeling. Thus, bile acid deconjugation is not unique to LLC but represents a reproducible component of the metabolic response to LLC colonization.

### LLC-Associated Remodeling of Fecal Bile Acid Conjugation Persists in a Complex Microbial Community

We next asked whether the bile acid conjugation phenotype observed in the gnotobiotic model was also detectable when LLC was administered to mice harboring a complex intestinal microbiota. In conventionally colonized mice, LLC treatment reduced several fecal bile acid measures relative to HBSS controls (**Figure 6A**). Total fecal bile acids were reduced following LLC treatment (Tukey-adjusted *P* = 0.0153), with particularly pronounced effects within the conjugated pool. Primary conjugated (*P* = 0.0003), secondary conjugated (*P* = 0.0129), and total conjugated bile acids (*P* = 0.0008) were all decreased, resulting in a marked reduction in the overall conjugated-to-unconjugated bile acid ratio (*P* = 0.0002). In contrast, total unconjugated bile acid abundance was comparatively preserved.

**Figure 6.**
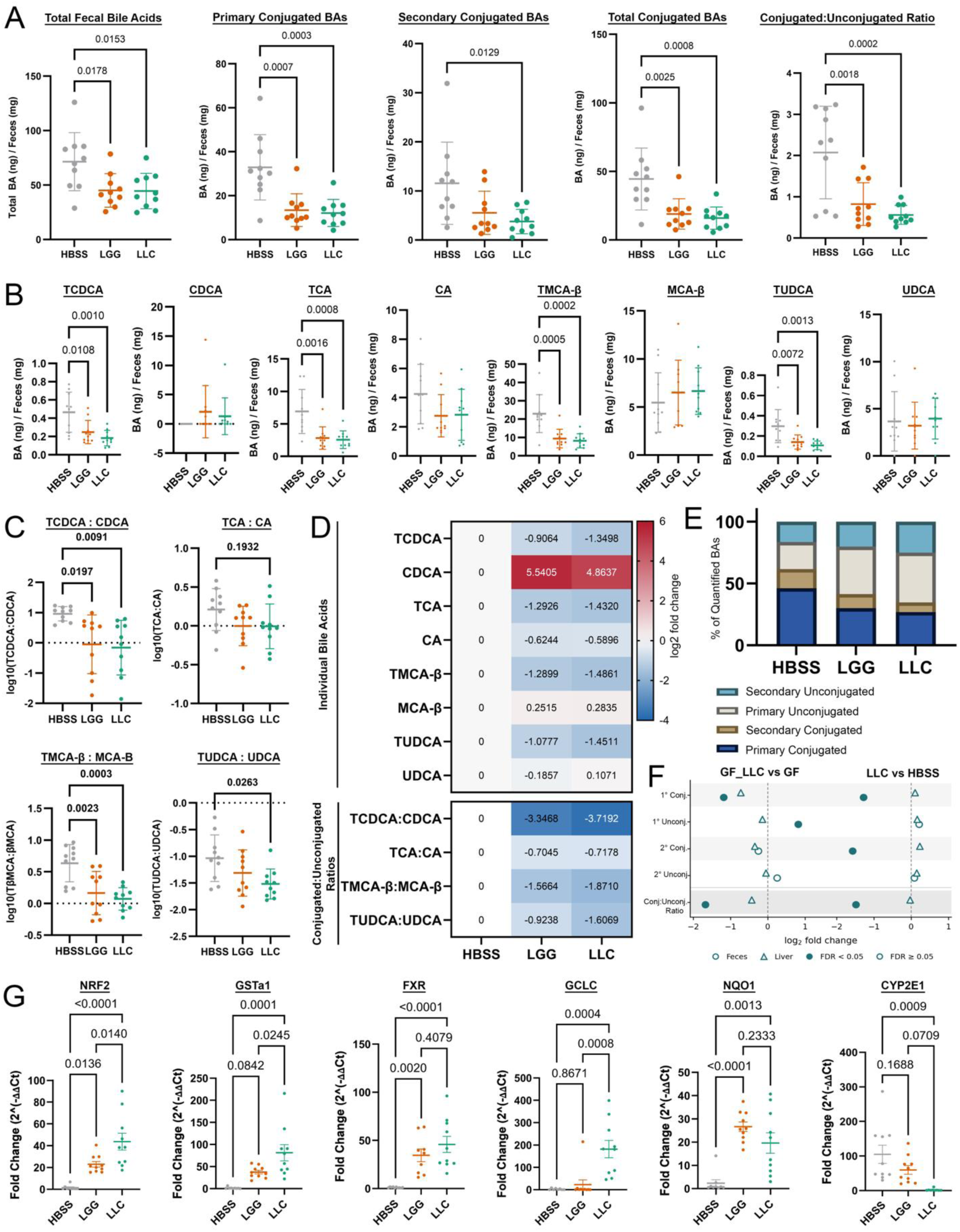
Probiotic treatment remodels the fecal bile acid pool and is associated with altered hepatic bile acid–responsive and cytoprotective gene expression in conventional mice. Conventional, standard-chow–fed mice were treated with HBSS, Lactococcus lactis subsp. cremoris (LLC), or Lactobacillus rhamnosus GG (LGG). **(A)** Total fecal bile acids, primary conjugated, secondary conjugated, and total conjugated bile acids, and the conjugated:unconjugated ratio. **(B)** Fecal concentrations of selected conjugated/unconjugated bile acid pairs: taurochenodeoxycholic acid (TCDCA)/chenodeoxycholic acid (CDCA), taurocholic acid (TCA)/cholic acid (CA), tauro-β-muricholic acid (TβMCA)/β-muricholic acid (βMCA), and tauroursodeoxycholic acid (TUDCA)/ursodeoxycholic acid (UDCA). **(C)** Log10-transformed TCDCA:CDCA, TCA:CA, TβMCA:βMCA, and TUDCA:UDCA ratios. **(D)** Heatmap summarizing individual bile acids and conjugated:unconjugated ratios in LGG- and LLC-treated mice relative to HBSS, displayed as log2 fold change. **(E)** Relative composition of the quantified fecal bile acid pool by primary conjugated, secondary conjugated, primary unconjugated, and secondary unconjugated classes. **(F)** Cross-compartment comparison of fecal and hepatic bile acid remodeling for GF_LLC versus GF and LLC versus HBSS, showing log2 fold changes for the four bile acid classes and conjugated:unconjugated ratio. Circles denote feces, triangles liver, filled symbols FDR-adjusted q < 0.05, and open symbols q ≥ 0.05; dashed lines indicate no change. **(G)** Hepatic RT-qPCR of Nrf2, Gsta1, Fxr, Gclc, Nqo1, and Cyp2e1. Points represent biological replicates; data are mean ± SD. Conventional groups contained n = 10 mice/group. Panels A–E and G were analyzed by one-way ANOVA with Tukey multiple-comparisons correction; exact adjusted p values are shown. Panel F used bile acid statistical models with Benjamini–Hochberg FDR correction.

LGG treatment produced a broadly similar pattern, including reductions in total fecal bile acids (*P* = 0.0178), primary conjugated bile acids (*P* = 0.0007), total conjugated bile acids (*P* = 0.0025), and the conjugated-to-unconjugated ratio (*P* = 0.0018) relative to HBSS controls. Thus, depletion of conjugated fecal bile acids was not unique to LLC. Nevertheless, the recurrence of this phenotype following LLC exposure in both monocolonized and conventionally colonized mice demonstrates that reduced fecal bile acid conjugation is a reproducible component of the LLC-associated metabolic response across distinct microbial contexts.

Analysis of individual bile acid species showed that the reduction in the conjugated pool reflected decreases across multiple taurine-conjugated bile acids (**Figure 6B**). Relative to HBSS controls, LLC treatment significantly reduced TCDCA (*P* = 0.0010), TCA (*P* = 0.0008), TβMCA (*P* = 0.0002), and TUDCA (*P* = 0.0013), whereas their corresponding unconjugated species exhibited smaller and more variable changes. Consistent with these species-level effects, the TCDCA:CDCA (*P* = 0.0091), TβMCA:βMCA (*P* = 0.0003), and TUDCA:UDCA (*P* = 0.0263) ratios were significantly reduced following LLC treatment (**Figure 6C**). In contrast, the TCA:CA ratio was not significantly altered (*P* = 0.1932), distinguishing the conventional response from the particularly pronounced TCA-to-CA shift observed following LLC monocolonization. The heatmap in **Figure 6D** summarizes these species-level changes and illustrates that the conventional LLC-associated phenotype was characterized primarily by depletion of conjugated fecal bile acids rather than uniform expansion of their unconjugated counterparts.

Relative bile acid composition demonstrated a similar pattern, with LLC-treated mice exhibiting a reduced proportional contribution of conjugated species and a corresponding greater contribution of unconjugated species to the quantified fecal bile acid pool (**Figure 6E**). These compositional data were interpreted alongside absolute concentrations because relative abundance alone cannot distinguish depletion of one bile acid class from expansion of another.

We next compared the magnitude of LLC-associated bile acid changes between fecal and hepatic compartments in both gnotobiotic and conventional settings (**Figure 6F**). In GF_LLC versus GF mice, fecal primary conjugated bile acids decreased to approximately 0.44-fold of GF levels (FDR-adjusted *q* = 0.025), primary unconjugated bile acids increased to approximately 1.76-fold (*q* = 0.025), and the conjugated-to-unconjugated ratio decreased to approximately 0.31-fold (*q* = 0.009). In contrast, none of the corresponding hepatic class-level measures differed significantly between GF_LLC and GF mice (all *q* ≈ 0.565).

A similar compartmental pattern was observed in conventionally colonized mice. LLC treatment reduced fecal primary conjugated bile acids to approximately 0.41-fold of HBSS levels (*q* < 0.001), secondary conjugated bile acids to approximately 0.34-fold (*q* = 0.0066), and the conjugated-to-unconjugated ratio to approximately 0.36-fold (*q* < 0.001), whereas corresponding hepatic bile acid classes remained comparatively unchanged (all *q* ≥ 0.749). Thus, across both microbial contexts, LLC-associated changes in bile acid conjugation were substantially more pronounced in feces than in liver, indicating compartmentalized remodeling of the bile acid pool rather than equivalent changes across intestinal and hepatic compartments.

Finally, we examined hepatic transcriptional pathways related to antioxidant defense, xenobiotic metabolism, and bile acid-associated signaling (**Figure 6G**). Relative to HBSS controls, LLC significantly increased hepatic expression of *Nrf2* (*P* < 0.0001), *Gsta1* (*P* = 0.0001), *Fxr* (*P* < 0.0001), *Gclc* (*P* = 0.0004), and *Nqo1* (*P* = 0.0013), while reducing *Cyp2e1* expression (*P* = 0.0009). These transcriptional changes occurred alongside the predominantly intestinal bile acid phenotype but do not establish a direct causal relationship between bile acid remodeling and hepatic gene regulation.

To determine whether probiotic-derived factors could elicit related transcriptional responses in a human hepatic cell model, HepG2 cells were exposed to bacterial culture supernatants derived from LLC or LGG. Supernatants from both strains significantly increased *GCLC* expression (*P* < 0.05), whereas *CYP2E1* expression was not significantly altered (**Supplementary Figure 5**). These findings indicate that soluble factors produced by LLC and LGG can induce antioxidant-associated transcriptional responses in a human hepatocyte-derived cell line, while regulation of xenobiotic-metabolizing genes such as *CYP2E1* may be dependent on the experimental context.

Together, these findings demonstrate that LLC reproducibly reduces fecal bile acid conjugation across both reductionist and complex microbial settings while hepatic class-level bile acid pools remain comparatively stable. This intestinal phenotype occurs alongside hepatic transcriptional changes involving antioxidant defense, xenobiotic metabolism, and *Fxr* expression. Considered together with the untargeted metabolomic and network analyses, these results identify altered intestinal bile acid conjugation as one component of a broader LLC-associated metabolic response spanning intestinal and hepatic compartments.

## Discussion

In this study, we demonstrate that *Lactococcus lactis* subsp. *cremoris* (LLC) exerts broad effects on hepatic metabolism that are associated with increased resistance to chemically induced liver injury. By integrating untargeted metabolomics across conventional and gnotobiotic models, functional injury experiments, network-based analysis, targeted bile acid profiling and hepatic transcriptional measurements, we identify a multidimensional LLC-associated metabolic response encompassing xenobiotic metabolism, mitochondrial and lipid-associated pathways, intestinal bile acid conjugation, and hepatic antioxidant signaling. Importantly, the bile acid analysis revealed that LLC does not uniformly remodel bile acids throughout the enterohepatic system. Instead, its most pronounced effects occur within the intestinal compartment, where LLC reproducibly reduced fecal conjugated bile acids while hepatic bile acid pools remain comparatively buffered. These findings position altered bile acid conjugation as one compartment of a broader LLC-associated metabolic program rather than a singular mechanism of hepatoprotection.

Under Western-style diet conditions, LLC induced substantial remodeling of both the circulating and hepatic metabolomes. The separation of LLC-treated mice from HBSS controls in serum and liver was accompanied by enrichment of circulating lipid associated pathways and hepatic pathways related to tryptophan metabolism and cytochrome P450 mediated xenobiotic processing. These observations extend previous work demonstrating systemic metabolic effects of LLC ^13, 14^ and are particularly relevant in the context of the liver, where xenobiotic metabolism and oxidative stress responses are tightly coupled ^8, 11^. Consistent with these metabolic signatures, LLC reduced hepatic injury in two mechanistically distinct models. Both LLC and LGG attenuated ethanol-induced lipid accumulation, whereas protection from APAP-associated centrilobular necrosis was more apparent following LLC treatment. Thus, LLC-associated hepatoprotection contains both effects shared with another lactic acid bacterium and responses that appear more strain-selective. This distinction is important, as the biological effects of probiotics are unlikely to be explained by a single metabolic property shared across strains ^37, 38^.

The gnotobiotic experiments provided a reductionist framework for distinguishing metabolic effects associated with LLC from those arising from microbial colonization. Monocolonization with LLC was sufficient to substantially alter the hepatic metabolome relative to germ-free controls, demonstrating that these responses do not require a complex microbial community. At the same time, LLC and LGG produced partially overlapping metabolic phenotypes, while conventionalization with a complete SPF microbiota generated a considerably broader response.

Network analysis further emphasized that the hepatic response to LLC is distributed across multiple coordinated metabolic programs rather than dominated by a single pathway. LLC-associated modules contained metabolite features mapping to xenobiotic metabolism, lipid oxidation, mitochondrial energy metabolism, and bile acid metabolism. Of these, drug metabolism–cytochrome P450 showed significant pathway enrichment, reinforcing the prominence of xenobiotic-associated metabolism observed in the broader metabolomic analyses.

Other LLC-associated modules contained top-ranked annotations related to electron transport, lipid oxidation, and bile acid biosynthesis, although these pathway associations did not reach conventional thresholds for statistical significance. Accordingly, these signals were interpreted as hypothesis-generating rather than evidence of definitive pathway enrichment. The appearance of bile acid biosynthesis among the top annotations of an LLC-associated module was nevertheless biologically notable given the direct interface between microbial enzymatic activity, bile acid chemistry, and host metabolism ^27, 39, 40^, and provided a rationale for targeted evaluation of bile acid metabolism.

This network-level observation was further supported by qualitative bile salt hydrolase assays demonstrating that LLC possesses bile acid deconjugation capacity in vitro. Both LLC and LGG cultures produced visible precipitation in the presence of conjugated bile acids, consistent with generation of less-soluble unconjugated bile acids. Precipitation was qualitatively more pronounced in LLC cultures than in LGG cultures under the conditions tested. Because this assay was qualitative and did not directly quantify substrate consumption, product generation, or enzyme kinetics, these observations should not be interpreted as demonstrating quantitatively greater BSH activity in LLC. Although LGG has previously characterized bile salt hydrolase activity ^36, 41^, LLC, a strain most commonly associated with dairy fermentation ^42^, has not, to our knowledge, been characterized for this function. These findings therefore identify bile acid deconjugation as a previously underappreciated metabolic property of LLC and provide a biological basis for examining whether this activity is reflected in vivo.

Targeted profiling revealed that microbial colonization had a particularly strong effect on fecal bile acid conjugation. Germ-free mice exhibited a fecal pool dominated by conjugated bile acids, consistent with the absence of microbial bile salt hydrolase activity and other microbial bile acid transformations ^22, 26, 27^. LLC monocolonization was sufficient to shift this profile toward a less conjugated state. The most striking molecular example was the reciprocal response of TCA and CA with TCA decreasing and CA increasing following LLC monocolonization. A directionally concordant pattern was observed for the murine TβMCA–βMCA pair, and these individual changes occurred together with a broader reduction in the total conjugated to unconjugated bile acid ratio.

Comparison with GF_SPF mice further illustrates the distinction between bile acid deconjugation and the broader repertoire of microbial bile acid metabolism. Conventionalization produced substantially greater expansion of unconjugated bile acids, including secondary species, than either monocolonizing strain. This is consistent with the requirement for multiple microbial enzymatic activities to generate the diversity of bile acid transformations present in a complex community ^25, 27, 39^. Thus, LLC alone can reproduce an important early step in microbial bile acid metabolism, deconjugation, but does not recapitulate the full range of downstream transformations generated by an intact microbiota.

The reduction in bile acid conjugation observed in gnotobiotic mice was also detectable when LLC was administered in the presence of a conventional microbiota. In conventional mice, LLC reduced primary conjugate, secondary conjugated, and total conjugated fecal bile acids, producing a substantial decline in the conjugated to unconjugated ratio. Importantly, total unconjugated bile acid abundance was comparatively preserved. The decrease in total fecal bile acids following LLC treatment therefore appears to be driven predominantly by depletion of conjugated species rather than by a reciprocal expansion of unconjugated or secondary bile acids.

At the individual metabolite level, conventional LLC treatment reduced several taurine-conjugated bile acids, including TCA, TβMCA, TCDCA, and TUDCA. However, the specific conjugated-unconjugated relationships differed between microbial contexts. Whereas TCA and CA exhibited a pronounced reciprocal response following LLC monocolonization, the TCA:CA ratio was not significantly altered by LLC in conventional mice. Other conjugated-to-unconjugated pairs showed stronger responses in the conventional setting. This context dependence likely reflects the substantially greater metabolic complexity of an intact intestinal community, in which LLC-derived activities occur alongside competing deconjugation, transformation, absorption, and cross-feeding processes ^27, 40^. The reproducible feature across models is therefore not alteration of a single bile acid pair, but a broader reduction in fecal bile acid conjugation.

LGG produced several directionally similar effects in conventional animals, including reductions in conjugated bile acid pools. These findings are consistent with bile acid deconjugation being a microbial function shared to varying degrees among lactic acid bacteria ^36, 41^. The significance of the LLC phenotype lies not in its uniqueness but in its reproducibility across reductionist and conventional microbial contexts and its occurrence alongside broader LLC-associated changes in hepatic metabolism. This distinction also highlights the importance of comparator strains when assigning functional properties to candidate probiotics ^37, 38^.

A central finding of the bile acid analysis was the marked compartmentalization of these effects. Direct comparison of fecal and hepatic bile acid changes demonstrated that LLC-associated alterations were consistently larger in feces than in liver. In GF_LLC mice, significant changes in fecal primary conjugated and primary unconjugated bile acids and in the conjugated-to-unconjugated ratio occurred without corresponding significant class-level changes in the liver. The same pattern was evident in conventional animals, where substantial depletion of fecal conjugated bile acids was not accompanied by parallel alterations in hepatic bile acid subclasses. These data indicate that the intestinal bile acid pool is substantially more responsive to LLC than the hepatic pool and suggest that hepatic bile acid composition can remain comparatively stable despite pronounced changes in the intestinal compartment.

The intestinal bile acid phenotype occurred alongside pronounced changes in hepatic pathways involved in antioxidant defense, xenobiotic metabolism, and bile acid-responsive signaling. In conventional mice, LLC increased hepatic expression of NRF2-responsive genes, including *Gsta1*, *Gclc*, and *Nqo1*, while reducing *Cyp2e1* expression. These transcriptional changes are consistent with the xenobiotic- and redox-associated pathways identified by untargeted metabolomics and WGCNA and provide a plausible biological context for the observed reduction in APAP-associated liver injury ^11^. CYP2E1 contributes directly to APAP bioactivation and generation of the reactive metabolite NAPQI ^30, 32^, whereas NRF2 coordinates multiple antioxidant and detoxification responses ^8, 43^. The convergence of metabolomic, transcriptional, and injury data therefore supports coordinated remodeling of hepatic stress-response and xenobiotic-metabolism pathways following LLC treatment.

LLC treatment also increased hepatic *Fxr* transcript abundance in conventional mice. Previous studies have demonstrated that microbial bile acid metabolism and FXR signaling can influence host lipid and energy metabolism, with increased bile salt hydrolase activity associated with altered bile acid reabsorption, depletion of selected conjugated bile acid species, modulation of intestinal FXR signaling, and protection from diet-induced weight gain ^20, 26, 44–46^. Although increased *Fxr* expression does not establish increased FXR activity, its occurrence alongside reduced fecal bile acid conjugation raises the possibility that bile acid-responsive signaling represents one component of the broader host response to LLC. Establishing such a relationship will require direct measurements of receptor activity and experimental manipulation of the relevant bile acid pathways.

To determine whether probiotic-derived factors could similarly influence antioxidant and xenobiotic-associated transcription in a human hepatic cell model, HepG2 cells were exposed to bacterial culture supernatants derived from LLC or LGG. Supernatants from both strains significantly increased *GCLC* expression (*P* < 0.05), whereas neither produced a significant change in *CYP2E1* expression. These findings indicate that soluble factors produced by LLC and LGG can induce antioxidant-associated transcriptional responses in a human hepatocyte-derived cell model, while also demonstrating that regulation of xenobiotic-metabolizing enzymes such as CYP2E1 is context dependent.

Other metabolite-sensitive pathways may also contribute to the LLC response. In particular, the recurring enrichment of tryptophan and xenobiotic metabolism raises the possibility that signaling through receptors such as AHR intersects with NRF2, FXR or other hepatic stress response pathways ^23, 47^. Because AHR activity was not measured directly, this remains a hypothesis for future investigation. More broadly, however, the diversity of metabolic pathways associated with LLC argues against a model in which its hepatoprotective effects arise from one metabolite class or receptor pathway.

Taken together, our findings support a model in which LLC modifies the gut–liver axis through several interconnected but not yet causally ordered processes. LLC is capable of altering bile acid conjugation within the intestinal compartment and is simultaneously associated with broader changes in hepatic xenobiotic, mitochondrial, and redox-related metabolism. Hepatic NRF2-associated transcription is increased and *Cyp2e1* expression reduced, providing a potential biological basis for increased resistance to oxidative and xenobiotic stress. Bile acid remodeling may contribute to these hepatic responses; however, the present experiments do not establish bile acids as an intermediary between LLC and hepatoprotection. Rather, altered intestinal bile acid conjugation represents one measurable component of a broader systems-level response to LLC.

Several limitations should be considered when interpreting these findings. Future studies incorporating tandem mass spectrometry, higher-confidence spectral annotation, and validation with authentic standards will be required to establish chemical identities of features not validated. Second, targeted bile acid profiling measured steady-state concentrations rather than metabolic flux and therefore cannot distinguish changes in microbial transformation from altered intestinal absorption, hepatic synthesis, biliary secretion, or enterohepatic cycling ^21, 48^. Likewise, although LLC monocolonization reduced fecal bile acid conjugation and LLC exhibited deconjugation capacity in vitro, this phenotype was not unique to LLC, and the qualitative precipitation assay does not establish reaction kinetics, substrate specificity, or the quantitative contribution of LLC-derived BSH activity in vivo ^36, 41^. Isotope-tracing studies, quantitative enzymatic assays, and direct measurements of bile acid transport and synthesis will be required to resolve these mechanisms.

Finally, the present experiments do not establish that bile acid remodeling mediates LLC-associated hepatic transcriptional responses or protection from liver injury. Although reduced fecal bile acid conjugation coincided with increased hepatic NRF2-associated genes and *Fxr* and reduced *Cyp2e1*, receptor activity was not directly measured, and increased *Fxr* transcript abundance should not be interpreted as evidence of increased FXR activity, which is governed by ligand availability and downstream receptor signaling ^15, 20^. Establishing causality will require genetic or pharmacologic perturbation of bile acid receptors, targeted manipulation of bile acid pools, or use of BSH-deficient bacterial strains. In addition, the relatively small gnotobiotic group sizes limited statistical power for some comparisons. Replication in larger cohorts and extension to additional dietary conditions, injury models, sexes, and human systems will be important for establishing the generalizability of these findings.

More broadly, these findings illustrate the value of combining complex and reductionist microbial models to resolve the metabolic effects of individual commensal strains. The gnotobiotic experiments establish that LLC alone is sufficient to alter both hepatic and metabolic organization and intestinal bile acid conjugation, whereas the conventional experiments demonstrate which features remain detectable within a complex microbiota. This approach also reveals an important distinction between functions that are reproducibly associated with LLC and those that are uniquely attributable to LLC. Defining these relationships will be critical for developing microbiome-based interventions in which strain selection is guided by measurable metabolic functions rather than taxonomic identity alone which may not often reflect the magnitude of phenotypic impact ^37, 38^.

## Materials and Methods

### Experimental Model and Subject Details Mice

All animal studies were approved by the Institutional Animal Care and Use Committee (IACUC) at Emory University and conducted in accordance with NIH guidelines. All experiments were conducted on C57BL/6J adult male mice. Conventional adult C57BL/6J mice were obtained from The Jackson Laboratory (Bar Harbor, ME) and maintained under standard barrier conditions with ad libitum access to food and water. Mice were housed under specific pathogen-free (SPF) conditions with support from the Emory Gnotobiotic Animal Core (EGAC). Germ-free (GF) mice were obtained directly from EGAC. For Western diet experiments, mice were fed a Western-style diet (high fat, high sucrose) beginning at 6–8 weeks of age and maintained on this diet for the duration of the study. For standard chow experiments, mice received standard laboratory chow. Specifically, mice were provided ad libitum access to sterilized 2019 Teklad Global 19% Protein Extruded Rodent Diet (Inotiv, Indianapolis, IN) or Western diet (D12079B, Research Diets, Inc.), as indicated, along with autoclaved drinking water. For select experiments involving probiotic administration to GF mice, animals were housed in hermetically sealed ISOcage P-Bioexclusion units (Tecniplast, West Chester, PA) within EGAC to maintain microbiological containment. All animals were maintained under standard environmental conditions, including a 12-hour light/dark cycle, and were monitored regularly for health and behavior. Where specified in the experimental design, mice received daily oral gavage of probiotics or vehicle control.

### Bacterial Culture

Lyophilized bacterial cultures were obtained from the American Type Culture Collection (ATCC). *Lactococcus lactis* subsp. *cremoris* (ATCC 19257) and *Lactobacillus rhamnosus* GG (ATCC 7469) were grown in MRS broth (de Man, Rogosa, and Sharpe; Millipore Sigma 1106610500) to stationary phase at 37°C. Cultures were then washed in sterile HBSS and administered by oral gavage once daily for the duration specified in the experimental design at a dose of 1 × 10⁸–10⁹ CFU total per dose. Vehicle-treated controls received HBSS alone.

### Cell Lines and Culture

HepG2 cells were maintained from stable freezer stocks and used at fewer than five passages. Cells were cultured in DMEM supplemented with 10% fetal bovine serum (FBS) and 1% penicillin/streptomycin at 37°C in a humidified incubator with 5% CO₂.

### Method Details Western Diet

To assess the global impact of probiotic exposure on the metabolome, C57BL/6 mice were randomly assigned to receive daily oral gavage of *Lactococcus lactis* subsp. *cremoris* (LLC), *Lactobacillus rhamnosus* GG (LGG), or vehicle control (HBSS) while concurrently maintained on a Western-style diet (D12079B, Research Diets, Inc.). This diet provides 17% of total caloric intake from protein, 43% from carbohydrates, and 40% from fat, reflecting a macronutrient composition associated with increased cardiometabolic risk. Probiotic administration and dietary intervention were initiated simultaneously and continued for four weeks. At the conclusion of the treatment period, mice were euthanized, and serum and tissues were harvested, flash-frozen in liquid nitrogen, and stored at −80°C until metabolomic analysis.

### Acetaminophen-Induced Liver Injury

To induce acute hepatotoxic injury, mice were fasted for 16 hours with free access to water prior to acetaminophen (APAP) administration. Acetaminophen was dissolved in HBSS containing 50% polyethylene glycol and administered by oral gavage at 300 mg/kg body weight, as specified in the experimental design. Animals were observed closely for signs of distress throughout the study period. Twenty-four hours following APAP administration, mice were euthanized and blood was collected by cardiac puncture for serum biochemical analyses. Livers were excised for histologic and biochemical assessment. For histopathologic evaluation, liver tissue was fixed in 10% neutral-buffered formalin, paraffin embedded, and sectioned. Hematoxylin and eosin (H&E) staining was performed using standard procedures. Sections were imaged at 20× magnification, and five non-overlapping fields per section were analyzed. The percentage of necrotic area was quantified in a blinded manner.

### Ethanol-Induced Hepatic Injury

Acute ethanol challenge was conducted using a single-binge model as previously described (Chen et al 2015, Saeedi et al 2020) ^11, 49^. Mice were fasted overnight (16 hours) with free access to water prior to ethanol administration. Ethanol was delivered by oral gavage at 6 g/kg body weight as a 70% ethanol solution diluted 70:30 (ethanol:water, v/v). Six hours after ethanol exposure, mice were euthanized and blood was collected as described above. Livers were harvested for histologic assessment of steatosis. For neutral lipid visualization, 7 μm-thick liver cryosections were fixed in formalin and stained with Oil Red O and counterstained with hematoxylin. Sections were imaged at 40× magnification, and representative fields were captured for analysis using ImageJ software.

### RNA Isolation and RT–qPCR

Liver tissue from conventional and germ-free C57BL/6 mice, with or without probiotic treatment (LLC or LGG, as specified in the experimental design), was mechanically homogenized in TRIzol reagent (Invitrogen, Carlsbad, CA) using a MagnaLyser instrument with MagnaLyser beads (Roche, Basel, Switzerland). Total RNA was extracted using the Aurum Total RNA Mini Kit (Bio-Rad, Hercules, CA) to ensure high purity and integrity. RNA concentration and quality were assessed spectrophotometrically prior to downstream analysis. Complementary DNA (cDNA) was synthesized from 1 µg of total RNA using the iScript cDNA Synthesis Kit (Bio-Rad) according to the manufacturer’s protocol. Relative transcript abundance was calculated using the comparative ΔΔCt method and normalized to the housekeeping gene 18S. Data are presented as fold change relative to the designated control group. The 18S ribosomal RNA control primers were: forward 5′-CGGAAAATAGCCTTCGCCATCAC-3′ and reverse 5′-ATCACTCGCTCCACCTCATCCT-3′. NRF2 primers were: forward 5′-CAGCATAGAGCAGGACATGGAG-3′ and reverse 5′-GAACAGCGGTAGTATCAGCCAG-3′. GCLC primers were: forward 5′-ACACCTGGATGATGCCAACGAG-3′ and reverse 5′-CCTCCATTGGTCGGAACTCTAC-3′. GSTA1 primers were: forward 5′-CTGCCTTGGCAAAAGATAGGACC-3′ and reverse 5′-CTTCCAGTAGGTGGATGTCCAC-3′. NQO1 primers were: forward 5′-GCCGAACACAAGAAGCTGGAAG-3′ and reverse 5′-GGCAAATCCTGCTACGAGCACT-3′. FXR (mouse Nr1h4) primers were: forward 5′-GGGATGAGTGTGAAGCCAGCTA-3′ and reverse 5′-GTGGCTGAACTTGAGGAAACGG-3′. CYP2E1 primers were: forward 5′-AGGCTGTCAAGGAGGTGCTACT-3′ and reverse 5′-AAAACCTCCGCACGTCCTTCCA-3′.

### Untargeted LCMS Metabolomics Sample Preparation

Liver tissue samples were extracted by adding 15 µL of ice-cold extraction solvent per milligram of tissue (33% LC–MS grade water [Thermo Scientific, 047146-K2] and 66% acetonitrile containing internal standards). Samples were homogenized using chilled MagNA Lyser Green Beads (Roche, REF 03358941001) in a MagNA Lyser instrument at maximum speed (7000 rpm) for 30 seconds. Serum samples were prepared by adding ice-cold acetonitrile at a 2:1 ratio (acetonitrile:serum, v/v). Following homogenization or solvent addition, samples were transferred to microcentrifuge tubes, vortexed briefly, and incubated on ice for 30 minutes to facilitate protein precipitation. Samples were then centrifuged at 14,000 × g for 10 minutes at 4°C. The clarified supernatants were transferred to autosampler vials and stored at −80°C until LC–MS analysis.

### HRM Instrument Analysis

Untargeted metabolomics was performed using high-resolution LCMS platforms. For western style diet fed and probiotic administered mice, LCMS instrument analysis was conducted as previously described in Gacasan et al 2026 ^14^. Samples were analyzed on a dual column and pump LC system (Thermo Ultimate3000 uHPLC) connected to a high resolution mass spectrometer (Thermo Scientific Q Exactive HF). The two analytical platforms consisted of a Waters Xbridge BEH Amide XP HILIC column (2.1 mm x 50 mm, 2.6 μm particle size) coupled with positive electrospray ionization (ESI+) and a Higgins Targa C18 column (2.1 mm x 50 mm, 3 μm) coupled with negative electrospray ionization (ESI-). The mobile phases included LCMS grade water (A), LCMS grade acetonitrile (B), and 2% formic acid in LCMS-grade water (C), and 10 mM ammonium acetate in LCMS grade water (D). For the HILIC chromatography gradient, an initial buffer ratio of 22.5% A, 75% B, and 2.5% C was held for 1.5-minutes before ascending to 75% A, 22.5% B, and 2.5% C for 4 minutes and ending with a 1-minute gradient hold. For the C18 chromatography gradient, an initial buffer ratio of 60 % A, 35% B, and 5% C was held for 1 minute before ascending to 0% A, 95% B, and 5% C for 3 minutes and ending with a 2-minute gradient hold. Column compartments were heated to 60 ℃. Flow rates were set to 0.35 mL/min for the first minute and 0.4 mL/min for the remaining 4 minutes. Mass spectra were collected at 120k resolution in a 85-1,275 *m/z* scan window.

For untargeted metabolomic samples acquired from GF mice, the LCMS platform consisted of a Thermo Scientific Vanquish Duo UHPLC and a Thermo Scientific Orbitrap ID-X. Two analytical platforms were used including hydrophilic interaction liquid chromatography (HILIC) coupled to positive electrospray ionization (ESI) and reversed phase C18 chromatography coupled to negative ESI. The HILIC method consisted of an Acquity BEH Amide HILIC column, 2.1 mm x 100 mm, 1.7 um (Waters, 186004801). Buffer A was water with 1 mM ammonium acetate and 0.1% formic acid and Buffer B was 95% acetonitrile with 1 mM ammonium acetate and 0.1% formic acid. For the HILIC gradient, an initial 0.5 min hold at 90% B was followed by a linear decrease to 20% B from 0.6 to 2.55 min, a 2 min hold, and then a 5-min re-equilibration period. The C18 method consisted of a Hypersil GOLD C18 column, 2.1 x 100 mm, 1.9 um (Thermo Scientific, 25003-102130). Buffer A was 0.1% formic acid in water and Buffer B was 99% acetonitrile with 0.1% formic acid. For the C18 gradient, an initial 0.5 min hold at 1% B was followed by a 0.75 min linear increase to 99% B, held for 3.75 min, and a 5-min re-equilibration period. Flow rates were 0.3 mL/min. The autosampler was set to 4 °Cand the column compartment at 45 °C. The mass spectrometer was operated at 120k resolution, and scans were collected for *m/z* 85-1,275. Tune parameters consisted of sheath gas at 50, auxiliary gas at 10, and sweep gas at 1. The spray voltage was set to 3.50 kV for ESI+ and -2.75 kV for ESI-. Ion transfer tube was set at 300 ℃ and vaporizer at 275 ℃.

### Untargeted Metabolomics Data Processing

Peak detection, noise removal, alignment and quantification were performed using a standardized workflow as previously described (Gacasan et al., 2026)^14^, with minor modifications as detailed: Adaptive processing for LCMS data (apLCMS) v6.3.3, with downstream quality control performed by xMSanalyzer v.2.0.8. Each metabolic feature was characterized by its m/z ratio, retention time, and peak intensity. For pathway enrichment analysis, differentially expressed features were normalized by log transformation, mean-centered and divided by the square root of the standard deviation of each variable and the top 10% of peaks used in mummichog v2.0 software. Spearman correlations and heatmaps were performed and prepared using Metaboanalyst v6.0.31. For volcano plots, metabolomics data were imported and reshaped using the tidyverse suite in R. Intensities were normalized to z-scores within each metabolite feature. Pairwise group comparisons were performed by calculating fold changes of mean z-scores and conducting two-sided t-tests for differential abundance. Features with log2 fold change > 1 and p < 0.05 were considered significant. Volcano plots highlighting enriched and depleted metabolites were generated using ggplot2. Detected features were putatively annotated using xMSannotator.

### Targeted Bile Acid Profiling

#### Target List and Spectral Library Construction

A targeted bile acid panel was constructed to include primary, secondary, and conjugated bile acids, with particular emphasis on amino acid–conjugated species in collaboration with the Emory Integrated Metabolomics and Lipidomics Core (EIMLC). Target compound information— including exact mass (m/z), chemical formula, and collision cross section (CCS) values—was curated from Stewart et al. (Analytical Chemistry, 2023) and incorporated into the spectral library. The EIMLC method included 23 bile acids for which authentic standards were available. These standards were used to confirm compound identity based on retention time (RT), m/z, and CCS alignment. Although glyco-ursodeoxycholic acid (GUDCA) was included among the available standards, it was not detected under the present chromatographic conditions and may co-elute with glycohyodeoxycholic acid (GHDCA).

#### Feature Identification and Confidence Scoring

Raw data were processed using MetaboScape (version 2025b). Extracted ion chromatograms (EICs) were manually reviewed for all candidate features to ensure appropriate peak integration and annotation quality. Alignment to the spectral library was evaluated using defined tolerances for m/z, retention time, MS/MS spectral similarity, mSigma, and CCS. Two acceptance windows were applied: m/z tolerance: 2–5 ppm, Retention time tolerance: 0.1–0.5 min, mSigma: 50–1000, MS/MS spectral match score: 400–800, CCS deviation: 1–5%. Each feature was assigned an MS/MS confidence score based on agreement with these parameters. Features falling outside tolerance limits were assigned a score of 0. Matches meeting upper-range criteria were assigned a score of 1, and those meeting stricter lower-range criteria were assigned a score of 2. Confidence scores were reported alongside quantitative outputs. Authentic standards were used to validate bile acids when RT, m/z, and CCS matched the reference library. Additional conjugated bile acids were tentatively annotated based on concordant m/z and CCS values.

#### Standard Curve Construction and Quantification

For bile acids with authentic standards, calibration curves were generated using serial dilutions. To preserve linearity, the three highest concentration points, where detector saturation or signal plateauing was observed were excluded unless otherwise specified. For UDCA, HDCA, and CDCA, which exhibited reduced linearity at higher concentrations, the five calibration points demonstrating the strongest linear relationship between concentration and peak area were selected. Eighteen bile acids were quantified absolutely and reported in ng per mg of liver tissue.

#### Data Curation and Quality Control

Features were excluded from downstream analysis under the following conditions: (1) the maximum sample intensity was lower than the mean blank intensity (e.g., GCDCA); (2) the ratio of maximum sample intensity to mean blank intensity was <5; or (3) manual review identified inadequate peak integration, chromatographic interference, or failure to meet predefined identification criteria. When duplicate features were determined to represent the same bile acid species on the basis of retention time and MS/MS similarity, their intensities were summed prior to downstream analysis. Signals exceeding the upper limit of the corresponding calibration curve were retained as estimated concentrations for comparative analyses but were flagged as outside the validated quantitative range and for potential re-injection. All extracted ion chromatogram (EIC) peaks were manually reviewed to confirm appropriate integration and, where possible, distinguish isomeric species using chromatographic separation and MS/MS fragmentation patterns.

Zero concentration values were not automatically excluded from analysis. For analyses performed on the original concentration scale, measured zero values were retained as zero. For analyses requiring logarithmic transformation, including log-transformed bile acid ratios and fold-change calculations, a bile acid–specific pseudocount equal to one-half of the minimum positive value observed for that analyte within the relevant dataset was added prior to transformation to permit inclusion of zero-valued observations. Conjugated-to-unconjugated and individual conjugated:unconjugated bile acid ratios were calculated at the individual-animal level prior to transformation and statistical testing. Values excluded on the basis of quality-control criteria were treated as missing rather than as zero and were omitted only from analyses involving that bile acid and any derived measures dependent upon it. Accordingly, the excluded GF_LLC βMCA measurement was treated as missing and reduced the sample size to *n* = 4 only for βMCA and βMCA-derived endpoints.

#### Semi-Quantitative Normalization

Bile acids lacking validated calibration curves were reported as semi-quantitative values. These were normalized to the peak area of α-muricholic acid (MCA-α), which served as an internal reference. Normalized peak areas were multiplied by the known concentration of MCA-α in each sample to express values in ng per mg of liver tissue.

#### Multivariate and Statistical Analysis of Untargeted Metabolomics

All statistical analyses and visualizations were performed in R (version 4.3.3) within the RStudio integrated development environment. Normalized feature intensity matrices were log-transformed and scaled (mean-centered and Pareto-scaled where indicated) prior to multivariate modeling to reduce heteroscedasticity and emphasize biologically meaningful variation. Principal component analysis (PCA) was performed using the prcomp function in base R and visualized using ggplot2 (v3.x). PCA score plots displayed individual samples as points colored by treatment group, with 95% confidence ellipses generated using stat_ellipse() to illustrate within-group dispersion. Scree plots were generated to display the proportion of variance explained by each principal component. Partial least squares–discriminant analysis (PLS-DA) was conducted using the mixOmics package. Model performance was evaluated using R² and Q² statistics obtained through cross-validation, and permutation testing was performed to assess model robustness and minimize overfitting. Variable importance in projection (VIP) scores were extracted and visualized using ranked bar plots generated with ggplot2.

Permutational multivariate analysis of variance (PERMANOVA) was performed using the vegan package (adonis2 function) based on Bray–Curtis dissimilarity matrices generated with vegdist. A total of 999 permutations were used to assess statistical significance. Pairwise PERMANOVA comparisons were conducted where indicated, and false discovery rate (FDR) correction was applied using the Benjamini–Hochberg method. Distance-based ordinations, including principal coordinates analysis (PCoA), were performed using vegan and visualized with ggplot2. Pairwise PERMANOVA results were summarized in formatted tables and displayed as heatmaps using ComplexHeatmap or pheatmap where appropriate. All plots were generated using ggplot2, with figure composition and multi-panel layouts arranged using patchwork.

#### Pathway Enrichment Analysis (Mummichog)

Metabolic pathway enrichment analysis was performed using mummichog v2.0. Significant features from pairwise comparisons were input into mummichog using KEGG pathway libraries.

Enrichment factor was calculated as:

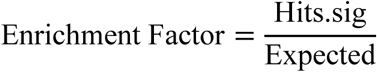

Significance was determined using Fisher’s exact test and gamma-adjusted p-values.

Directional overlap across contrasts was summarized using UpSet plots, which provide a quantitative visualization of intersections among multiple pathway sets. For each pairwise contrast, significantly enriched pathways were split into two direction-labeled sets (e.g., *higher in LLC* vs *higher in HBSS/GF*), generating “signed” pathway lists that preserve biological directionality. These signed pathway sets were then converted into a binary membership matrix (pathways × contrasts), where each entry indicated whether a given pathway was present in a specific direction-labeled set. UpSet visualizations were generated in R (RStudio) using ComplexUpset (built on ComplexHeatmap), with intersection bars representing the number of pathways shared across specific combinations of contrasts. The matrix layout beneath the bars indicated membership of each pathway in each contrast-specific set, enabling rapid identification of pathways that recur across multiple comparisons with consistent direction of change. In addition to signed overlap, “unsigned” UpSet plots were also generated by collapsing direction labels to assess recurrence of pathway identity irrespective of direction, and membership tables were exported to transparently report pathway composition for each intersection.

#### Weighted Gene Correlation Network Analysis (WGCNA)

WGCNA was conducted as previously described in Gacasan et al 2026^14^. Median-summarized and m/z-calibrated untargeted metabolomics data were processed in R (v4.3.3) using the WGCNA package (v1.72-5). Features were annotated using a combined m/z and retention time identifier (mz rt), log₂-transformed, and filtered using the goodSamplesGenes function to remove low-quality samples and low-variance features prior to network construction. Sample clustering was first performed using hierarchical clustering to evaluate overall sample relationships and identify potential outliers. The soft-thresholding power (β = 11) was selected using the pickSoftThreshold function to approximate scale-free network topology while preserving network connectivity. Pairwise Pearson correlations between metabolites were computed and transformed into an adjacency matrix, which was subsequently converted into a Topological Overlap Matrix (TOM) to enhance network robustness by incorporating shared neighbor relationships.

Metabolites were clustered using average linkage hierarchical clustering based on 1 – TOM dissimilarity. Network modules were identified using dynamic tree cutting (deepSplit = 2) with minimum cluster size parameters of 100, 250, and 500 to evaluate module granularity. A minimum cluster size of 250 was selected for downstream analyses to balance module resolution and biological interpretability. Module eigengenes (MEs), representing the first principal component of each module, were correlated with experimental traits, including probiotic treatment and microbial status. Module–trait associations were assessed using Pearson correlation, and statistical significance was determined using Student’s t-test. Module assignments, eigengene values, module–trait correlation matrices, and annotated metabolite feature lists were exported for downstream pathway enrichment and biological interpretation.

#### Bile Salt Hydrolase (BSH) Assays

LLC and LGG were cultured in MRS medium supplemented with either a mixed bile acid pool (0.5% w/v glycocholic acid, glycodeoxycholic acid, taurocholic acid, and taurodeoxycholic acid) or individual bile acids supplied at 1 mM. Bile acid deconjugation activity was assessed qualitatively by the appearance of visible precipitation, reflecting the reduced solubility of unconjugated bile acids following enzymatic deconjugation.

### Statistical Analysis

#### Statistics

Statistical analysis was performed using Prism 10 (GraphPad, San Diego, CA). Significance was defined as \**P* ≤.05, \*\**P* ≤.01, \*\*\**P* ≤.001, \*\*\*\**P* ≤.0001. Data presented as mean +/-SD unless otherwise noted.

## Data Availability Statement

Raw metabolomics data generated in this study are available through Metabolomics Workbench (study DOIs pending). Untargeted metabolomics feature tables and additional datasets are available via the Emory Dataverse.

## Code Availability

All computational code used for microbiome profiling, metabolomics analysis, and other workflows are publicly available via Zenodo at 10.5281/zenodo.22772601. This repository includes scripts for statistical modeling, data visualization, and multiomic integration, including correlation and network based approaches as well as feature tables, and targeted and untargeted datasets.

All other relevant data supporting the findings of this study are available from the corresponding author upon reasonable request.

## Author Contributions

CAG and RMJ conceived and designed the experiments. Mouse experiments were conducted by CAG and GMW with the assistance of LCA, MEB and the staff at the Emory Gnotobiotic Animal Core. CAG conducted sample preparation for untargeted metabolomics. Untargeted mass spectrometry data acquisition was performed by JW and DPJ. Targeted bile acid analysis including sample preparation was conducted by the Emory Lipidomics Core. All downstream HRMS analysis was performed by CAG. GMW conducted cell culture experiments and subsequent RT-qPCR analysis. All other analyses and experiments were conducted and analyzed by CAG and GMW. CAG and RMJ wrote the manuscript.

## Acknowledgments

The Emory Integrated Cellular Imaging Core (RRID:SCR_023534) and the Emory Cancer Tissue and Pathology Shared Resource are subsidized by the Emory University School of Medicine. The Emory Gnotobiotic Animal Core especially Caroline Addis and Amanda Metzger for their assistance with animal handling as well as Sean D. Kelly from the Sampson Lab for dissection assistance.

This study was funded by a NRSA F30 training grant from the National Institute of Diabetes and Digestive and Kidney Diseases *F30DK139762* (CAG), and *F30DK134204* (LCA), and *K12HD072245* (MEB). The funder played no role in study design, data collection, analysis and interpretation of data, or the writing of this manuscript. This content is solely the responsibility of the authors and does not necessarily represent the official view of the National Institutes of Health or Centers for Disease Control and Prevention.

## Declaration of generative AI and AI-assisted technologies in the manuscript preparation process

During the preparation of this work, the author(s) used ChatGPT and Claude for code debugging and proofreading purposes. After using this tool/service, the author(s) reviewed and edited the content as needed and take(s) full responsibility for the content of the published article.

