## Supplementary for "*Lactococcus lactis* subsp. cremoris Reprograms the Gut-Liver Axis and Protects Against Liver Injury"

**SUPPLEMENTARY DATA**

C. Anthony Gacasan[1], Jaclyn Weinberg[2], Gabrielle M. Webster[1], Crystal R. Naudin[1], ViLinh Tran[2], Lauren C. Askew[1], Maria E. Barbian[3], Dean P. Jones[2], and Rheinallt M. Jones[1*]

Affiliations:

[1]Division of Gastroenterology, Hepatology, and Nutrition, Department of Pediatrics, [2]Division of Pulmonary, Allergy, Critical Care and Sleep Medicine, Department of Medicine, [3]Division of Neonatology, Department of Pediatrics, Emory University School of Medicine, Atlanta GA, 30322.

**Corresponding author:** * Rheinallt M. Jones, Division of Gastroenterology, Hepatology, and Nutrition, Department of Pediatrics, Emory University School of Medicine, 615 Michael Street, Atlanta GA, 30322, Tel: (404) 727-7231, Fax: (404) 727-8538

***Disclosures: The authors have no conflicts of interest.***

***Key words:*** Probiotics, Microbiome, Gut-Liver Axis, Xenobiotic Metabolism, FXR, NRF2, CYP450, Bile Acids, Metabolomics, Acetaminophen, Alcohol, Western Diet

**Running head:**

Supported by the National Institutes of Health grants F30DK134204 (LCA), F30DK139762 (CAG), and K12HD072245 (MEB).

A

| Serum | F.Model | R2 | pval | p.adj |
|---|---|---|---|---|
| HBSS vs LGG | 2.7933 | 0.14112 | 0.047 | 0.0705 |
| HBSS vs LLC | 9.6994 | 0.36328 | 0.001 | 0.003 |
| LGG vs LLC | 2.7137 | 0.13101 | 0.104 | 0.104 |

B

| Liver | F.Model | R2 | pval | p.adj |
|---|---|---|---|---|
| HBSS vs LGG | 3.0584 | 0.15248 | 0.092 | 0.092 |
| HBSS vs LLC | 9.4025 | 0.35612 | 0.001 | 0.003 |
| LGG vs LLC | 4.5474 | 0.20168 | 0.009 | 0.0135 |

Supplementary Table 1. Pairwise PERMANOVA analysis of untargeted metabolomics profiles from conventionally raised, Western-style diet–fed mice (n = 10 per group). (A) Serum; (B) Liver.

A

| 7 group PERMANOVA - Liver | Df | SumOfSqs | R2 | F | Pr(>F) |
|---|---|---|---|---|---|
| Group | 6 | 182250.1 | 0.267895 | 2.561472 | 1.00E-04 |
| Residual | 42 | 498053.9 | 0.732105 | NA | NA |
| Total | 48 | 680304 | 1 | NA | NA |

B

| Group1 | Group2 | F | R2 | p | p_adj_fdr |
|---|---|---|---|---|---|
| HBSS | GF | 3.68114 | 0.220677 | 0.0003 | 0.0014 |
| HBSS | GF_SPF | 3.397742 | 0.207208 | 0.0002 | 0.0014 |
| HBSS | GF_LLC | 2.94676 | 0.184787 | 0.0004 | 0.0014 |
| LLC | GF | 4.53563 | 0.258652 | 0.0003 | 0.0014 |
| LLC | GF_SPF | 3.725157 | 0.222728 | 0.0002 | 0.0014 |
| LLC | GF_LLC | 3.690976 | 0.221136 | 0.0004 | 0.0014 |
| LGG | GF | 2.831651 | 0.17886 | 0.0009 | 0.0027 |
| HBSS | LLC | 1.641811 | 0.083588 | 0.0012 | 0.002864 |
| HBSS | GF_LGG | 2.916717 | 0.195533 | 0.0015 | 0.002864 |
| LLC | GF_LGG | 3.64206 | 0.232838 | 0.0015 | 0.002864 |
| LGG | GF_SPF | 2.384833 | 0.155012 | 0.0014 | 0.002864 |
| LGG | GF_LLC | 2.389479 | 0.155267 | 0.0026 | 0.00455 |
| GF_SPF | GF_LLC | 2.497411 | 0.237907 | 0.0065 | 0.0105 |
| GF | GF_SPF | 2.222785 | 0.217434 | 0.0076 | 0.010631 |
| GF | GF_LLC | 2.85821 | 0.26323 | 0.0081 | 0.010631 |
| GF_SPF | GF_LGG | 2.440569 | 0.258519 | 0.008 | 0.010631 |
| GF | GF_LGG | 2.75567 | 0.282469 | 0.009 | 0.011118 |
| LGG | GF_LGG | 2.285131 | 0.159966 | 0.0109 | 0.012717 |
| LLC | LGG | 1.20086 | 0.062542 | 0.1125 | 0.124342 |
| HBSS | LGG | 1.158182 | 0.060454 | 0.183 | 0.19215 |
| GF_LLC | GF_LGG | 1.002974 | 0.125325 | 0.4607 | 0.4607 |

Supplementary Table 2. PERMANOVA of untargeted liver metabolomic profiles from standard chow–fed conventional (n = 10 per group) and germ-free/gnotobiotic (n = 5 per group) mice. (A) Overall analysis including all groups (LLC, LGG, HBSS, GF_LLC, GF_LGG, GF, GF_SPF). (B) Pairwise comparisons across groups.

| Pathway_unsigned | PathwayName | n_comparisons_unsigned | Comparisons_present_unsigned | total_features_across_sets | max_n_features_in_any_set | max_n_ec_in_any_set | directions_observed | n_directions |
|---|---|---|---|---|---|---|---|---|
| Methionine and cysteine metabolism | Methionine and cysteine metabolism | 4 | GF_LLC/HBSS; LLC/GF; LLC/GF_LLC; LLC/HBSS | 54 | 36 | 17 | GF_LLC; HBSS; LLC | 3 |
| Purine metabolism | Purine metabolism | 4 | GF_LLC/HBSS; LLC/GF; LLC/GF_LLC; LLC/HBSS | 52 | 33 | 21 | GF_LLC; HBSS; LLC | 3 |
| Aspartate and asparagine metabolism | Aspartate and asparagine metabolism | 3 | GF_LLC/HBSS; LLC/GF; LLC/GF_LLC | 114 | 74 | 43 | GF_LLC; LLC | 2 |
| Tryptophan metabolism | Tryptophan metabolism | 3 | GF_LLC/HBSS; LLC/GF; LLC/GF_LLC | 103 | 70 | 46 | GF_LLC; HBSS; LLC | 3 |
| Tyrosine metabolism | Tyrosine metabolism | 3 | GF_LLC/HBSS; LLC/GF; LLC/GF_LLC | 105 | 67 | 55 | GF_LLC; HBSS; LLC | 3 |
| Urea cycle/amino group metabolism | Urea cycle/amino group metabolism | 3 | GF_LLC/HBSS; LLC/GF; LLC/GF_LLC | 103 | 66 | 43 | GF_LLC; LLC | 2 |
| Glycerophospholipid metabolism | Glycerophospholipid metabolism | 3 | GF_LLC/HBSS; LLC/GF; LLC/GF_LLC | 52 | 38 | 23 | GF_LLC; LLC | 2 |
| Glycine, serine, alanine and threonine metabolism | Glycine, serine, alanine and threonine metabolism | 3 | GF_LLC/HBSS; LLC/GF; LLC/GF_LLC | 53 | 38 | 21 | GF_LLC; LLC | 2 |
| Arachidonic acid metabolism | Arachidonic acid metabolism | 3 | GF_LLC/HBSS; LLC/GF; LLC/GF_LLC | 55 | 34 | 18 | GF_LLC; LLC | 2 |
| Arginine and Proline Metabolism | Arginine and Proline Metabolism | 3 | GF_LLC/HBSS; LLC/GF; LLC/GF_LLC | 52 | 34 | 21 | GF_LLC; LLC | 2 |
| Pyrimidine metabolism | Pyrimidine metabolism | 3 | GF_LLC/HBSS; LLC/GF; LLC/GF_LLC | 46 | 33 | 21 | GF_LLC; LLC | 2 |
| Drug metabolism - cytochrome P450 | Drug metabolism - cytochrome P450 | 3 | GF_LLC/HBSS; LLC/GF; LLC/GF_LLC | 40 | 27 | 21 | GF_LLC; HBSS; LLC | 3 |
| Histidine metabolism | Histidine metabolism | 3 | GF_LLC/HBSS; LLC/GF; LLC/GF_LLC | 33 | 27 | 16 | GF; GF_LLC; HBSS | 3 |
| C21-steroid hormone biosynthesis and metabolism | C21-steroid hormone biosynthesis and metabolism | 3 | GF_LLC/HBSS; LLC/GF; LLC/GF_LLC | 38 | 22 | 14 | GF_LLC; LLC | 2 |
| Xenobiotics metabolism | Xenobiotics metabolism | 3 | GF_LLC/HBSS; LLC/GF; LLC/GF_LLC | 34 | 22 | 10 | GF; HBSS; LLC | 3 |
| Vitamin B3 (nicotinate and nicotinamide) metabolism | Vitamin B3 (nicotinate and nicotinamide) metabolism | 3 | GF_LLC/HBSS; LLC/GF; LLC/GF_LLC | 29 | 20 | 11 | GF_LLC; HBSS; LLC | 3 |
| Androgen and estrogen biosynthesis and metabolism | Androgen and estrogen biosynthesis and metabolism | 3 | GF_LLC/HBSS; LLC/GF; LLC/GF_LLC | 25 | 19 | 9 | GF; GF_LLC; LLC | 3 |
| Aminosugars metabolism | Aminosugars metabolism | 3 | GF_LLC/HBSS; LLC/GF; LLC/GF_LLC | 30 | 18 | 12 | GF_LLC; LLC | 2 |
| Valine, leucine and isoleucine degradation | Valine, leucine and isoleucine degradation | 3 | GF_LLC/HBSS; LLC/GF; LLC/GF_LLC | 30 | 16 | 12 | GF_LLC; LLC | 2 |
| Alanine and Aspartate Metabolism | Alanine and Aspartate Metabolism | 3 | GF_LLC/HBSS; LLC/GF; LLC/GF_LLC | 24 | 15 | 10 | GF_LLC; LLC | 2 |
| Butanoate metabolism | Butanoate metabolism | 3 | GF_LLC/HBSS; LLC/GF; LLC/GF_LLC | 28 | 15 | 15 | GF_LLC; LLC | 2 |
| Bile acid biosynthesis | Bile acid biosynthesis | 3 | GF_LLC/HBSS; LLC/GF; LLC/GF_LLC | 23 | 14 | 8 | GF; HBSS; LLC | 3 |
| Biopterin metabolism | Biopterin metabolism | 3 | GF_LLC/HBSS; LLC/GF; LLC/GF_LLC | 20 | 14 | 9 | GF_LLC; HBSS; LLC | 3 |
| De novo fatty acid biosynthesis | De novo fatty acid biosynthesis | 3 | GF_LLC/HBSS; LLC/GF; LLC/GF_LLC | 23 | 14 | 11 | GF_LLC; LLC | 2 |
| Omega-3 fatty acid metabolism | Omega-3 fatty acid metabolism | 3 | GF_LLC/HBSS; LLC/GF; LLC/GF_LLC | 24 | 12 | 7 | GF; HBSS; LLC | 3 |
| Vitamin B6 (pyridoxine) metabolism | Vitamin B6 (pyridoxine) metabolism | 3 | GF_LLC/HBSS; LLC/GF; LLC/GF_LLC | 20 | 12 | 8 | GF_LLC; LLC | 2 |
| Fatty acid activation | Fatty acid activation | 3 | GF_LLC/HBSS; LLC/GF; LLC/GF_LLC | 17 | 9 | 8 | GF; GF_LLC; LLC | 3 |
| Vitamin B5 - CoA biosynthesis from pantothenate | Vitamin B5 - CoA biosynthesis from pantothenate | 3 | GF_LLC/HBSS; LLC/GF; LLC/GF_LLC | 14 | 7 | 3 | GF_LLC; LLC | 2 |
| CoA Catabolism | CoA Catabolism | 3 | GF_LLC/HBSS; LLC/GF; LLC/GF_LLC | 12 | 6 | 2 | GF_LLC; LLC | 2 |
| N-Glycan biosynthesis | N-Glycan biosynthesis | 3 | GF_LLC/HBSS; LLC/GF; LLC/HBSS | 10 | 6 | 4 | HBSS; LLC | 2 |
| Pentose phosphate pathway | Pentose phosphate pathway | 3 | GF_LLC/HBSS; LLC/GF; LLC/GF_LLC | 12 | 6 | 3 | HBSS; LLC | 2 |
| Linoleate metabolism | Linoleate metabolism | 2 | LLC/GF; LLC/GF_LLC | 68 | 48 | 29 | GF; LLC | 2 |
| Prostaglandin formation from arachidonate | Prostaglandin formation from arachidonate | 2 | GF_LLC/HBSS; LLC/GF | 43 | 31 | 16 | GF_LLC; LLC | 2 |
| Lysine metabolism | Lysine metabolism | 2 | GF_LLC/HBSS; LLC/GF | 39 | 30 | 15 | GF_LLC; LLC | 2 |
| Leukotriene metabolism | Leukotriene metabolism | 2 | GF_LLC/HBSS; LLC/GF | 42 | 29 | 16 | GF; GF_LLC | 2 |
| Glutathione Metabolism | Glutathione Metabolism | 2 | GF_LLC/HBSS; LLC/GF | 22 | 19 | 9 | GF; HBSS | 2 |
| Vitamin A (retinol) metabolism | Vitamin A (retinol) metabolism | 2 | GF_LLC/HBSS; LLC/GF | 30 | 19 | 9 | GF_LLC; LLC | 2 |
| Glutamate metabolism | Glutamate metabolism | 2 | LLC/GF; LLC/GF_LLC | 26 | 18 | 8 | GF; GF_LLC | 2 |
| Sialic acid metabolism | Sialic acid metabolism | 2 | GF_LLC/HBSS; LLC/GF | 16 | 14 | 9 | GF_LLC; LLC | 2 |
| Carnitine shuttle | Carnitine shuttle | 2 | LLC/GF; LLC/GF_LLC | 15 | 13 | 9 | LLC | 1 |
| Vitamin H (biotin) metabolism | Vitamin H (biotin) metabolism | 2 | GF_LLC/HBSS; LLC/GF | 16 | 11 | 6 | HBSS; LLC | 2 |
| Phosphatidylinositol phosphate metabolism | Phosphatidylinositol phosphate metabolism | 2 | GF_LLC/HBSS; LLC/GF | 10 | 8 | 6 | HBSS; LLC | 2 |
| Porphyrin metabolism | Porphyrin metabolism | 2 | GF_LLC/HBSS; LLC/GF | 11 | 8 | 6 | GF_LLC; LLC | 2 |
| Vitamin E metabolism | Vitamin E metabolism | 2 | GF_LLC/HBSS; LLC/GF | 12 | 8 | 8 | GF; HBSS | 2 |
| Fructose and mannose metabolism | Fructose and mannose metabolism | 2 | LLC/GF; LLC/GF_LLC | 9 | 6 | 4 | LLC | 1 |
| Keratan sulfate degradation | Keratan sulfate degradation | 2 | GF_LLC/HBSS; LLC/GF | 7 | 5 | 4 | HBSS; LLC | 2 |
| Fatty Acid Metabolism | Fatty Acid Metabolism | 2 | LLC/GF; LLC/GF_LLC | 7 | 4 | 4 | GF; LLC | 2 |
| Glycolysis and Gluconeogenesis | Glycolysis and Gluconeogenesis | 2 | GF_LLC/HBSS; LLC/GF | 7 | 4 | 4 | HBSS; LLC | 2 |
| Hyaluronan Metabolism | Hyaluronan Metabolism | 2 | GF_LLC/HBSS; LLC/GF | 6 | 4 | 3 | HBSS; LLC | 2 |
| Limonene and pinene degradation | Limonene and pinene degradation | 2 | LLC/GF; LLC/GF_LLC | 7 | 4 | 3 | GF; LLC | 2 |
| Omega-6 fatty acid metabolism | Omega-6 fatty acid metabolism | 2 | GF_LLC/HBSS; LLC/GF | 6 | 4 | 2 | GF_LLC; LLC | 2 |
| Squalene and cholesterol biosynthesis | Squalene and cholesterol biosynthesis | 2 | GF_LLC/HBSS; LLC/GF | 8 | 4 | 4 | GF; HBSS | 2 |
| Alkaloid biosynthesis II | Alkaloid biosynthesis II | 2 | GF_LLC/HBSS; LLC/GF | 5 | 3 | 2 | HBSS; LLC | 2 |
| Propanoate metabolism | Propanoate metabolism | 2 | GF_LLC/HBSS; LLC/GF | 6 | 3 | 3 | GF_LLC; LLC | 2 |
| Vitamin B1 (thiamin) metabolism | Vitamin B1 (thiamin) metabolism | 2 | GF_LLC/HBSS; LLC/GF | 5 | 3 | 2 | HBSS; LLC | 2 |

**Supplementary Table S3. Recurrently enriched metabolic pathways identified across pairwise comparisons independent of directional assignment.**

This table summarizes metabolic pathways identified by mummichog that recur across multiple pairwise contrasts spanning germ-free, monocolonized, and conventional contexts. Pathways are collapsed to their unsigned form (i.e., direction of change not considered), allowing identification of metabolic programs that are repeatedly perturbed regardless of whether they are increased or decreased in LLC-containing states. For each pathway, the number of comparisons in which it was enriched is reported, along with robustness metrics including feature counts and enzyme commission (EC) coverage. This table provides a global overview of recurrent metabolic remodeling independent of probiotic-associated directionality.

| PathwayName | Pathway_signed | Direction | n_comparisons_unsigned | Direction_consistency | n_directions | median_t_summary | n_features | n_ec |
|---|---|---|---|---|---|---|---|---|
| Androgen and estrogen biosynthesis and metabolism | Androgen and estrogen biosynthesis and metabolism_higher_in_GF | GF | 3 | flipped | 3 | -3.8956 | 19 | 9 |
| Bile acid biosynthesis | Bile acid biosynthesis_higher_in_GF | GF | 3 | flipped | 3 | -3.5738 | 14 | 8 |
| Fatty Acid Metabolism | Fatty Acid Metabolism_higher_in_GF | GF | 2 | flipped | 2 | -4.908 | 3 | 3 |
| Fatty acid activation | Fatty acid activation_higher_in_GF | GF | 3 | flipped | 3 | -3.9977 | 9 | 8 |
| Glutamate metabolism | Glutamate metabolism_higher_in_GF | GF | 2 | flipped | 2 | -0.22725 | 18 | 8 |
| Glutathione Metabolism | Glutathione Metabolism_higher_in_GF | GF | 2 | flipped | 2 | -4.0925 | 19 | 9 |
| Histidine metabolism | Histidine metabolism_higher_in_GF | GF | 3 | flipped | 3 | -4.1939 | 27 | 16 |
| Leukotriene metabolism | Leukotriene metabolism_higher_in_GF | GF | 2 | flipped | 2 | -3.3661 | 29 | 16 |
| Limonene and pinene degradation | Limonene and pinene degradation_higher_in_GF | GF | 2 | flipped | 2 | -5.4267 | 4 | 3 |
| Linoleate metabolism | Linoleate metabolism_higher_in_GF | GF | 2 | flipped | 2 | -4.4676 | 48 | 29 |
| Omega-3 fatty acid metabolism | Omega-3 fatty acid metabolism_higher_in_GF | GF | 3 | flipped | 3 | -4.2268 | 12 | 7 |
| Squalene and cholesterol biosynthesis | Squalene and cholesterol biosynthesis_higher_in_GF | GF | 2 | flipped | 2 | -2.54965 | 4 | 4 |
| Vitamin E metabolism | Vitamin E metabolism_higher_in_GF | GF | 2 | flipped | 2 | -4.32525 | 8 | 8 |
| Xenobiotics metabolism | Xenobiotics metabolism_higher_in_GF | GF | 3 | flipped | 3 | -5.07725 | 22 | 10 |
| Alanine and Aspartate Metabolism | Alanine and Aspartate Metabolism_higher_in_GF_LLC | GF_LLC | 3 | flipped | 2 | -1.62083 | 5 | 3 |
| Aminosugars metabolism | Aminosugars metabolism_higher_in_GF_LLC | GF_LLC | 3 | flipped | 2 | -0.38495 | 6 | 4 |
| Androgen and estrogen biosynthesis and metabolism | Androgen and estrogen biosynthesis and metabolism_higher_in_GF_LLC | GF_LLC | 3 | flipped | 3 | 3.99455 | 4 | 2 |
| Arachidonic acid metabolism | Arachidonic acid metabolism_higher_in_GF_LLC | GF_LLC | 3 | flipped | 2 | -2.6077 | 13 | 8 |
| Arginine and Proline Metabolism | Arginine and Proline Metabolism_higher_in_GF_LLC | GF_LLC | 3 | flipped | 2 | -0.31102 | 10 | 7 |
| Aspartate and asparagine metabolism | Aspartate and asparagine metabolism_higher_in_GF_LLC | GF_LLC | 3 | flipped | 2 | -0.82205 | 22 | 15 |
| Biopterin metabolism | Biopterin metabolism_higher_in_GF_LLC | GF_LLC | 3 | flipped | 3 | -8.61235 | 2 | 2 |
| Butanoate metabolism | Butanoate metabolism_higher_in_GF_LLC | GF_LLC | 3 | flipped | 2 | -1.14148 | 9 | 7 |
| C21-steroid hormone biosynthesis and metabolism | C21-steroid hormone biosynthesis and metabolism_higher_in_GF_LLC | GF_LLC | 3 | flipped | 2 | 4.9877 | 12 | 7 |
| CoA Catabolism | CoA Catabolism_higher_in_GF_LLC | GF_LLC | 3 | flipped | 2 | 4.68045 | 6 | 2 |
| De novo fatty acid biosynthesis | De novo fatty acid biosynthesis_higher_in_GF_LLC | GF_LLC | 3 | flipped | 2 | 5.0166 | 5 | 3 |
| Drug metabolism - cytochrome P450 | Drug metabolism - cytochrome P450_higher_in_GF_LLC | GF_LLC | 3 | flipped | 3 | -10.0745 | 7 | 7 |
| Fatty acid activation | Fatty acid activation_higher_in_GF_LLC | GF_LLC | 3 | flipped | 3 | 5.0166 | 3 | 2 |
| Glutamate metabolism | Glutamate metabolism_higher_in_GF_LLC | GF_LLC | 2 | flipped | 2 | -9.05065 | 8 | 2 |
| Glycerophospholipid metabolism | Glycerophospholipid metabolism_higher_in_GF_LLC | GF_LLC | 3 | flipped | 2 | 0.594925 | 8 | 4 |
| Glycine, serine, alanine and threonine metabolism | Glycine, serine, alanine and threonine metabolism_higher_in_GF_LLC | GF_LLC | 3 | flipped | 2 | -0.67367 | 10 | 4 |
| Histidine metabolism | Histidine metabolism_higher_in_GF_LLC | GF_LLC | 3 | flipped | 3 | -7.9251 | 3 | 2 |
| Leukotriene metabolism | Leukotriene metabolism_higher_in_GF_LLC | GF_LLC | 2 | flipped | 2 | 3.3563 | 13 | 7 |
| Lysine metabolism | Lysine metabolism_higher_in_GF_LLC | GF_LLC | 2 | flipped | 2 | 6.5702 | 9 | 5 |
| Methionine and cysteine metabolism | Methionine and cysteine metabolism_higher_in_GF_LLC | GF_LLC | 4 | flipped | 3 | -1.82293 | 9 | 3 |
| Omega-6 fatty acid metabolism | Omega-6 fatty acid metabolism_higher_in_GF_LLC | GF_LLC | 2 | flipped | 2 | 4.1652 | 4 | 2 |
| Porphyrin metabolism | Porphyrin metabolism_higher_in_GF_LLC | GF_LLC | 2 | flipped | 2 | 5.1692 | 3 | 3 |
| Propanoate metabolism | Propanoate metabolism_higher_in_GF_LLC | GF_LLC | 2 | flipped | 2 | 5.0166 | 3 | 2 |
| Prostaglandin formation from arachidonate | Prostaglandin formation from arachidonate_higher_in_GF_LLC | GF_LLC | 2 | flipped | 2 | 3.6959 | 12 | 6 |
| Purine metabolism | Purine metabolism_higher_in_GF_LLC | GF_LLC | 4 | flipped | 3 | -5.6409 | 4 | 3 |
| Pyrimidine metabolism | Pyrimidine metabolism_higher_in_GF_LLC | GF_LLC | 3 | flipped | 2 | -0.19225 | 7 | 6 |
| Sialic acid metabolism | Sialic acid metabolism_higher_in_GF_LLC | GF_LLC | 2 | flipped | 2 | 0.0287 | 2 | 2 |
| Tryptophan metabolism | Tryptophan metabolism_higher_in_GF_LLC | GF_LLC | 3 | flipped | 3 | -9.05245 | 12 | 10 |
| Tyrosine metabolism | Tyrosine metabolism_higher_in_GF_LLC | GF_LLC | 3 | flipped | 3 | -9.05065 | 17 | 10 |
| Urea cycle/amino group metabolism | Urea cycle/amino group metabolism_higher_in_GF_LLC | GF_LLC | 3 | flipped | 2 | -0.6309 | 21 | 12 |
| Valine, leucine and isoleucine degradation | Valine, leucine and isoleucine degradation_higher_in_GF_LLC | GF_LLC | 3 | flipped | 2 | -1.09883 | 8 | 6 |
| Vitamin A (retinol) metabolism | Vitamin A (retinol) metabolism_higher_in_GF_LLC | GF_LLC | 2 | flipped | 2 | 5.22905 | 11 | 7 |
| Vitamin B3 (nicotinate and nicotinamide) metabolism | Vitamin B3 (nicotinate and nicotinamide) metabolism_higher_in_GF_LLC | GF_LLC | 3 | flipped | 3 | -7.3483 | 3 | 2 |
| Vitamin B5 - CoA biosynthesis from pantothenate | Vitamin B5 - CoA biosynthesis from pantothenate_higher_in_GF_LLC | GF_LLC | 3 | flipped | 2 | 4.7069 | 7 | 3 |
| Vitamin B6 (pyridoxine) metabolism | Vitamin B6 (pyridoxine) metabolism_higher_in_GF_LLC | GF_LLC | 3 | flipped | 2 | 2.122925 | 6 | 5 |

**Supplementary Table S4. Direction-aware metabolic pathways associated with LLC across colonization contexts.** This table presents directionally annotated metabolic pathways identified by mummichog across pairwise comparisons spanning germ-free, monocolonized, and conventional conditions. Each pathway is reported in signed form (e.g., “higher in LLC” or “higher in comparator”), enabling interpretation of probiotic-associated increases and decreases separately. For each pathway, the table includes the dominant direction of change, directional consistency across contrasts, median test statistics, and pathway robustness metrics (number of matched metabolite features and enzyme commission (EC) coverage). By incorporating directionality, this table distinguishes conserved LLC-associated metabolic programs from background shifts attributable to colonization state alone.

| Pathway_signed | GF vs GF_LLC | GF vs LLC | GF_LLC vs HBSS | HBSS vs LLC |
|---|---|---|---|---|
| 3-oxo-10R-octadecatrienoate beta-oxidation_higher_in_LLC | FALSE | TRUE | FALSE | FALSE |
| Alanine and Aspartate Metabolism_higher_in_GF_LLC | TRUE | FALSE | FALSE | FALSE |
| Alanine and Aspartate Metabolism_higher_in_HBSS | FALSE | FALSE | TRUE | FALSE |
| Alanine and Aspartate Metabolism_higher_in_LLC | FALSE | TRUE | FALSE | FALSE |
| Alkaloid biosynthesis II_higher_in_GF_LLC | FALSE | FALSE | TRUE | FALSE |
| Alkaloid biosynthesis II_higher_in_LLC | FALSE | TRUE | FALSE | FALSE |
| Aminosugars metabolism_higher_in_GF_LLC | TRUE | FALSE | FALSE | FALSE |
| Aminosugars metabolism_higher_in_HBSS | FALSE | FALSE | TRUE | FALSE |
| Aminosugars metabolism_higher_in_LLC | FALSE | TRUE | FALSE | FALSE |
| Androgen and estrogen biosynthesis and metabolism_higher_in_GF | TRUE | TRUE | FALSE | FALSE |
| Androgen and estrogen biosynthesis and metabolism_higher_in_HBSS | FALSE | FALSE | TRUE | FALSE |
| Arachidonic acid metabolism_higher_in_GF_LLC | TRUE | FALSE | FALSE | FALSE |
| Arachidonic acid metabolism_higher_in_HBSS | FALSE | FALSE | TRUE | FALSE |
| Arachidonic acid metabolism_higher_in_LLC | FALSE | TRUE | FALSE | FALSE |
| Arginine and Proline Metabolism_higher_in_GF_LLC | TRUE | FALSE | FALSE | FALSE |
| Arginine and Proline Metabolism_higher_in_HBSS | FALSE | FALSE | TRUE | FALSE |
| Arginine and Proline Metabolism_higher_in_LLC | FALSE | TRUE | FALSE | FALSE |
| Ascorbate (Vitamin C) and Aldarate Metabolism_higher_in_GF | FALSE | TRUE | FALSE | FALSE |
| Aspartate and asparagine metabolism_higher_in_GF_LLC | TRUE | FALSE | FALSE | FALSE |
| Aspartate and asparagine metabolism_higher_in_HBSS | FALSE | FALSE | TRUE | FALSE |
| Aspartate and asparagine metabolism_higher_in_LLC | FALSE | TRUE | FALSE | FALSE |
| Beta-Alanine metabolism_higher_in_LLC | FALSE | TRUE | FALSE | FALSE |
| Bile acid biosynthesis_higher_in_GF | TRUE | TRUE | FALSE | FALSE |
| Bile acid biosynthesis_higher_in_GF_LLC | FALSE | FALSE | TRUE | FALSE |
| Biopterin metabolism_higher_in_GF_LLC | TRUE | FALSE | TRUE | FALSE |
| Biopterin metabolism_higher_in_LLC | FALSE | TRUE | FALSE | FALSE |
| Blood Group Biosynthesis_higher_in_LLC | FALSE | TRUE | FALSE | FALSE |
| Butanoate metabolism_higher_in_GF_LLC | TRUE | FALSE | FALSE | FALSE |
| Butanoate metabolism_higher_in_HBSS | FALSE | FALSE | TRUE | FALSE |
| Butanoate metabolism_higher_in_LLC | FALSE | TRUE | FALSE | FALSE |
| C21-steroid hormone biosynthesis and metabolism_higher_in_GF | TRUE | FALSE | FALSE | FALSE |
| C21-steroid hormone biosynthesis and metabolism_higher_in_HBSS | FALSE | FALSE | TRUE | FALSE |
| C21-steroid hormone biosynthesis and metabolism_higher_in_LLC | FALSE | TRUE | FALSE | FALSE |
| Carbon fixation_higher_in_LLC | FALSE | TRUE | FALSE | FALSE |
| Carnitine shuttle_higher_in_GF | TRUE | FALSE | FALSE | FALSE |
| Carnitine shuttle_higher_in_LLC | FALSE | TRUE | FALSE | FALSE |
| Chondroitin sulfate degradation_higher_in_LLC | FALSE | TRUE | FALSE | FALSE |
| CoA Catabolism_higher_in_GF | TRUE | FALSE | FALSE | FALSE |
| CoA Catabolism_higher_in_HBSS | FALSE | FALSE | TRUE | FALSE |
| CoA Catabolism_higher_in_LLC | FALSE | TRUE | FALSE | FALSE |
| De novo fatty acid biosynthesis_higher_in_GF | TRUE | FALSE | FALSE | FALSE |
| De novo fatty acid biosynthesis_higher_in_HBSS | FALSE | FALSE | TRUE | FALSE |
| De novo fatty acid biosynthesis_higher_in_LLC | FALSE | TRUE | FALSE | FALSE |
| Di-unsaturated fatty acid beta-oxidation_higher_in_GF | TRUE | FALSE | FALSE | FALSE |
| Drug metabolism - cytochrome P450_higher_in_GF_LLC | TRUE | FALSE | TRUE | FALSE |
| Drug metabolism - cytochrome P450_higher_in_LLC | FALSE | TRUE | FALSE | FALSE |
| Drug metabolism - other enzymes_higher_in_LLC | FALSE | TRUE | FALSE | FALSE |
| Electron transport chain_higher_in_GF | FALSE | TRUE | FALSE | FALSE |
| Fatty Acid Metabolism_higher_in_GF | TRUE | TRUE | FALSE | FALSE |
| Fatty acid activation_higher_in_GF | TRUE | TRUE | FALSE | FALSE |
| Fatty acid activation_higher_in_HBSS | FALSE | FALSE | TRUE | FALSE |
| Fatty acid oxidation_higher_in_GF | TRUE | FALSE | FALSE | FALSE |
| Fructose and mannose metabolism_higher_in_GF | TRUE | FALSE | FALSE | FALSE |
| Fructose and mannose metabolism_higher_in_LLC | FALSE | TRUE | FALSE | FALSE |
| Galactose metabolism_higher_in_LLC | FALSE | TRUE | FALSE | FALSE |
| Glutamate metabolism_higher_in_GF | FALSE | TRUE | FALSE | FALSE |
| Glutamate metabolism_higher_in_GF_LLC | TRUE | FALSE | FALSE | FALSE |
| Glutathione Metabolism_higher_in_GF | FALSE | TRUE | FALSE | FALSE |
| Glutathione Metabolism_higher_in_GF_LLC | FALSE | FALSE | TRUE | FALSE |
| Glycerophospholipid metabolism_higher_in_GF_LLC | TRUE | FALSE | FALSE | FALSE |
| Glycerophospholipid metabolism_higher_in_HBSS | FALSE | FALSE | TRUE | FALSE |
| Glycerophospholipid metabolism_higher_in_LLC | FALSE | TRUE | FALSE | FALSE |
| Glycine, serine, alanine and threonine metabolism_higher_in_GF_LLC | TRUE | FALSE | FALSE | FALSE |
| Glycine, serine, alanine and threonine metabolism_higher_in_HBSS | FALSE | FALSE | TRUE | FALSE |
| Glycine, serine, alanine and threonine metabolism_higher_in_LLC | FALSE | TRUE | FALSE | FALSE |
| Glycolysis and Gluconeogenesis_higher_in_GF_LLC | FALSE | FALSE | TRUE | FALSE |
| Glycolysis and Gluconeogenesis_higher_in_LLC | FALSE | TRUE | FALSE | FALSE |
| Glycosphingolipid biosynthesis - ganglioseries_higher_in_LLC | FALSE | TRUE | FALSE | FALSE |
| Glycosphingolipid biosynthesis - globoseries_higher_in_LLC | FALSE | TRUE | FALSE | FALSE |
| Glycosphingolipid biosynthesis - lactoseries_higher_in_LLC | FALSE | TRUE | FALSE | FALSE |
| Glycosphingolipid biosynthesis - neolactoseries_higher_in_LLC | FALSE | TRUE | FALSE | FALSE |
| Glycosphingolipid metabolism_higher_in_LLC | FALSE | TRUE | FALSE | FALSE |
| Heparan sulfate degradation_higher_in_LLC | FALSE | TRUE | FALSE | FALSE |
| Hexose phosphorylation_higher_in_LLC | FALSE | TRUE | FALSE | FALSE |
| Histidine metabolism_higher_in_GF | FALSE | TRUE | FALSE | FALSE |
| Histidine metabolism_higher_in_GF_LLC | TRUE | FALSE | TRUE | FALSE |

| Pathway_signed | GF vs GF_LLC | GF vs LLC | GF_LLC vs HBSS | HBSS vs LLC |
|---|---|---|---|---|
| Keratan sulfate degradation_higher_in_LLC | FALSE | TRUE | FALSE | FALSE |
| Leukotriene metabolism_higher_in_GF | FALSE | TRUE | FALSE | FALSE |
| Leukotriene metabolism_higher_in_HBSS | FALSE | FALSE | TRUE | FALSE |
| Limonene and pinene degradation_higher_in_GF | TRUE | TRUE | FALSE | FALSE |
| Linoleate metabolism_higher_in_GF | TRUE | TRUE | FALSE | FALSE |
| Lysine metabolism_higher_in_HBSS | FALSE | FALSE | TRUE | FALSE |
| Lysine metabolism_higher_in_LLC | FALSE | TRUE | FALSE | FALSE |
| Methionine and cysteine metabolism_higher_in_GF_LLC | TRUE | FALSE | FALSE | FALSE |
| Methionine and cysteine metabolism_higher_in_HBSS | FALSE | FALSE | TRUE | TRUE |
| Methionine and cysteine metabolism_higher_in_LLC | FALSE | TRUE | FALSE | FALSE |
| Mono-unsaturated fatty acid beta-oxidation_higher_in_GF | TRUE | FALSE | FALSE | FALSE |
| N-Glycan Degradation_higher_in_LLC | FALSE | TRUE | FALSE | FALSE |
| N-Glycan biosynthesis_higher_in_GF_LLC | FALSE | FALSE | TRUE | FALSE |
| N-Glycan biosynthesis_higher_in_LLC | FALSE | TRUE | FALSE | TRUE |
| Nitrogen metabolism_higher_in_LLC | FALSE | TRUE | FALSE | FALSE |
| O-Glycan biosynthesis_higher_in_LLC | FALSE | TRUE | FALSE | FALSE |
| Omega-3 fatty acid metabolism_higher_in_GF | TRUE | TRUE | FALSE | FALSE |
| Omega-3 fatty acid metabolism_higher_in_GF_LLC | FALSE | FALSE | TRUE | FALSE |
| Omega-6 fatty acid metabolism_higher_in_HBSS | FALSE | FALSE | TRUE | FALSE |
| Omega-6 fatty acid metabolism_higher_in_LLC | FALSE | TRUE | FALSE | FALSE |
| Pentose and Glucuronate Interconversions_higher_in_LLC | FALSE | TRUE | FALSE | FALSE |
| Pentose phosphate pathway_higher_in_GF | TRUE | FALSE | FALSE | FALSE |
| Pentose phosphate pathway_higher_in_GF_LLC | FALSE | FALSE | TRUE | FALSE |
| Pentose phosphate pathway_higher_in_LLC | FALSE | TRUE | FALSE | FALSE |
| Phosphatidylinositol phosphate metabolism_higher_in_GF_LLC | FALSE | FALSE | TRUE | FALSE |
| Phosphatidylinositol phosphate metabolism_higher_in_LLC | FALSE | TRUE | FALSE | FALSE |
| Porphyrin metabolism_higher_in_HBSS | FALSE | FALSE | TRUE | FALSE |
| Porphyrin metabolism_higher_in_LLC | FALSE | TRUE | FALSE | FALSE |
| Propanoate metabolism_higher_in_HBSS | FALSE | FALSE | TRUE | FALSE |
| Propanoate metabolism_higher_in_LLC | FALSE | TRUE | FALSE | FALSE |
| Prostaglandin formation from arachidonate_higher_in_HBSS | FALSE | FALSE | TRUE | FALSE |
| Prostaglandin formation from arachidonate_higher_in_LLC | FALSE | TRUE | FALSE | FALSE |
| Prostaglandin formation from dihomo gama-linoleic acid_higher_in_LLC | FALSE | TRUE | FALSE | FALSE |
| Purine metabolism_higher_in_GF_LLC | TRUE | FALSE | TRUE | FALSE |
| Purine metabolism_higher_in_LLC | FALSE | TRUE | FALSE | TRUE |
| Putative anti-Inflammatory metabolites formation from EPA_higher_in_GF | FALSE | TRUE | FALSE | FALSE |
| Pyrimidine metabolism_higher_in_GF_LLC | TRUE | FALSE | FALSE | FALSE |
| Pyrimidine metabolism_higher_in_HBSS | FALSE | FALSE | TRUE | FALSE |
| Pyrimidine metabolism_higher_in_LLC | FALSE | TRUE | FALSE | FALSE |
| Pyruvate Metabolism_higher_in_GF | FALSE | TRUE | FALSE | FALSE |
| Saturated fatty acids beta-oxidation_higher_in_GF_LLC | TRUE | FALSE | FALSE | FALSE |
| Selenoamino acid metabolism_higher_in_LLC | FALSE | TRUE | FALSE | FALSE |
| Sialic acid metabolism_higher_in_HBSS | FALSE | FALSE | TRUE | FALSE |
| Sialic acid metabolism_higher_in_LLC | FALSE | TRUE | FALSE | FALSE |
| Squalene and cholesterol biosynthesis_higher_in_GF | FALSE | TRUE | FALSE | FALSE |
| Squalene and cholesterol biosynthesis_higher_in_GF_LLC | FALSE | FALSE | TRUE | FALSE |
| Starch and Sucrose Metabolism_higher_in_LLC | FALSE | TRUE | FALSE | FALSE |
| TCA cycle_higher_in_GF | FALSE | TRUE | FALSE | FALSE |
| Tryptophan metabolism_higher_in_GF_LLC | TRUE | FALSE | TRUE | FALSE |
| Tryptophan metabolism_higher_in_LLC | FALSE | TRUE | FALSE | FALSE |
| Tyrosine metabolism_higher_in_GF_LLC | TRUE | FALSE | TRUE | FALSE |
| Tyrosine metabolism_higher_in_LLC | FALSE | TRUE | FALSE | FALSE |
| Ubiquinone Biosynthesis_higher_in_LLC | FALSE | TRUE | FALSE | FALSE |
| Urea cycle/amino group metabolism_higher_in_GF_LLC | TRUE | FALSE | FALSE | FALSE |
| Urea cycle/amino group metabolism_higher_in_HBSS | FALSE | FALSE | TRUE | FALSE |
| Urea cycle/amino group metabolism_higher_in_LLC | FALSE | TRUE | FALSE | FALSE |
| Valine, leucine and isoleucine degradation_higher_in_GF_LLC | TRUE | FALSE | FALSE | FALSE |
| Valine, leucine and isoleucine degradation_higher_in_HBSS | FALSE | FALSE | TRUE | FALSE |
| Valine, leucine and isoleucine degradation_higher_in_LLC | FALSE | TRUE | FALSE | FALSE |
| Vitamin A (retinol) metabolism_higher_in_HBSS | FALSE | FALSE | TRUE | FALSE |
| Vitamin A (retinol) metabolism_higher_in_LLC | FALSE | TRUE | FALSE | FALSE |
| Vitamin B1 (thiamin) metabolism_higher_in_GF_LLC | FALSE | FALSE | TRUE | FALSE |
| Vitamin B1 (thiamin) metabolism_higher_in_LLC | FALSE | TRUE | FALSE | FALSE |
| Vitamin B2 (riboflavin) metabolism_higher_in_GF | FALSE | TRUE | FALSE | FALSE |
| Vitamin B3 (nicotinate and nicotinamide) metabolism_higher_in_GF_LLC | TRUE | FALSE | TRUE | FALSE |
| Vitamin B3 (nicotinate and nicotinamide) metabolism_higher_in_LLC | FALSE | TRUE | FALSE | FALSE |
| Vitamin B5 - CoA biosynthesis from pantothenate_higher_in_GF | TRUE | FALSE | FALSE | FALSE |
| Vitamin B5 - CoA biosynthesis from pantothenate_higher_in_HBSS | FALSE | FALSE | TRUE | FALSE |
| Vitamin B5 - CoA biosynthesis from pantothenate_higher_in_LLC | FALSE | TRUE | FALSE | FALSE |
| Vitamin B6 (pyridoxine) metabolism_higher_in_GF_LLC | TRUE | FALSE | FALSE | FALSE |
| Vitamin B6 (pyridoxine) metabolism_higher_in_HBSS | FALSE | FALSE | TRUE | FALSE |
| Vitamin B6 (pyridoxine) metabolism_higher_in_LLC | FALSE | TRUE | FALSE | FALSE |
| Vitamin B9 (folate) metabolism_higher_in_LLC | FALSE | TRUE | FALSE | FALSE |
| Vitamin D3 (cholecalciferol) metabolism_higher_in_GF | FALSE | TRUE | FALSE | FALSE |
| Vitamin E metabolism_higher_in_GF | FALSE | TRUE | FALSE | FALSE |
| Vitamin E metabolism_higher_in_GF_LLC | FALSE | FALSE | TRUE | FALSE |

**Supplementary Table S5. Complete direction-aware pathway membership matrix used for UpSet intersection analysis.**

This table contains the full boolean membership matrix for all directionally annotated pathways included in the UpSet analysis. Each column corresponds to one pairwise comparison (GF vs GF_LLC, GF vs LLC, GF_LLC vs HBSS, HBSS vs LLC), and entries indicate whether a given signed pathway was enriched in that contrast. This table provides full computational transparency and enables independent reconstruction of the direction-aware intersection analysis presented in Figure 3.

A

| RT (min) | CCS | Compound Name | Formula | Validated |
|---|---|---|---|---|
| 21.9 | 205.38743 | Dehydrolithocholic Acid-DHLCA | C24H38O3 | Yes |
| 21.9 | 202.820324 | Lithocholic Acid-LCA | C24H40O3 | Yes |
| 20.3 | 201.566797 | Hyodeoxycholic Acid-HDCA | C24H40O4 | Yes |
| 20.3 | 199.858695 | Ursodeoxycholic Acid-UDCA | C24H40O4 | Yes |
| 21.1 | 203.274907 | Chenodeoxycholic Acid-CDCA | C24H40O4 | Yes |
| 21.2 | 202.235188 | Deoxycholic Acid-DCA | C24H40O4 | Yes |
| 18.4 | 203.898137 | Muricholic Acid-MCA-alpha | C24H40O5 | Yes |
| 19.1 | 202.488939 | Muricholic Acid-MCA-beta | C24H40O5 | Yes |
| 20.2 | 202.859795 | Cholic Acid-CA | C24H40O5 | Yes |
| 18.1 | 201.078505 | Glycohyodeoxycholic Acid-GHDCA | C26H43NO5 | No |
| 18.1 | 200.930601 | Glycoursodeoxycholic Acid-GUDCA | C26H43NO5 | No |
| 20.3 | 200.486907 | Glycochenodeoxycholic Acid-GCDCA | C26H43NO5 | Yes |
| 18.1 | 201.908997 | Glycocholic Acid-GCA | C26H43NO6 | Yes |
| 21.3 | 206.343789 | Taurolithocholic Acid-TLCA | C26H45NO5S | Yes |
| 14 | 207.489119 | Tauroursodeoxycholic Acid-TUDCA | C26H45NO6S | Yes |
| 14.2 | 207.857782 | Taurohyodeoxycholic Acid-THDCA | C26H45NO6S | Yes |
| 18.8 | 206.973 | Taurochenodeoxycholic Acid-TCDCA | C26H45NO6S | Yes |
| 19.6 | 206.014496 | Taurodeoxycholic Acid-TDCA | C26H45NO6S | Yes |
| 6.3 | | Tauro-Muricholic Acid-TMCA-omega | C26H45NO7S | Yes |
| 6.8 | | Tauro-Muricholic Acid-TMCA-alpha | C26H45NO7S | Yes |
| 7.1 | | Tauro-Muricholic Acid-TMCA-beta | C26H45NO7S | Yes |
| 10.8 | | Tauro-Muricholic Acid-TMCA-gamma | C26H45NO7S | Yes |
| 14.8 | 207.317349 | Taurocholic Acid -TCA | C26H45NO7S | Yes |

B

| | Narrow (lower limit) | Wide (Upper limit) |
|---|---|---|
| m/z (ppm) | 2.0 | 5.0 |
| RT (min) | 0.1 | 0.5 |
| MSigma | 50 | 1000 |
| MS/MS score | 800 | 400 |
| CCS (%) | 1.0 | 5.0 |

**Supplementary Table S6. Fecal bile acid standards and alignment criteria. (A)** Comprehensive list of bile acid standards provided by the Emory Integrated Metabolomics and Lipidomics Core (EIMLC) used for targeted fecal bile acid quantification. **(B)** Alignment tolerances and scoring criteria applied for metabolite matching, including mass accuracy and retention time thresholds used to ensure confident feature annotation.

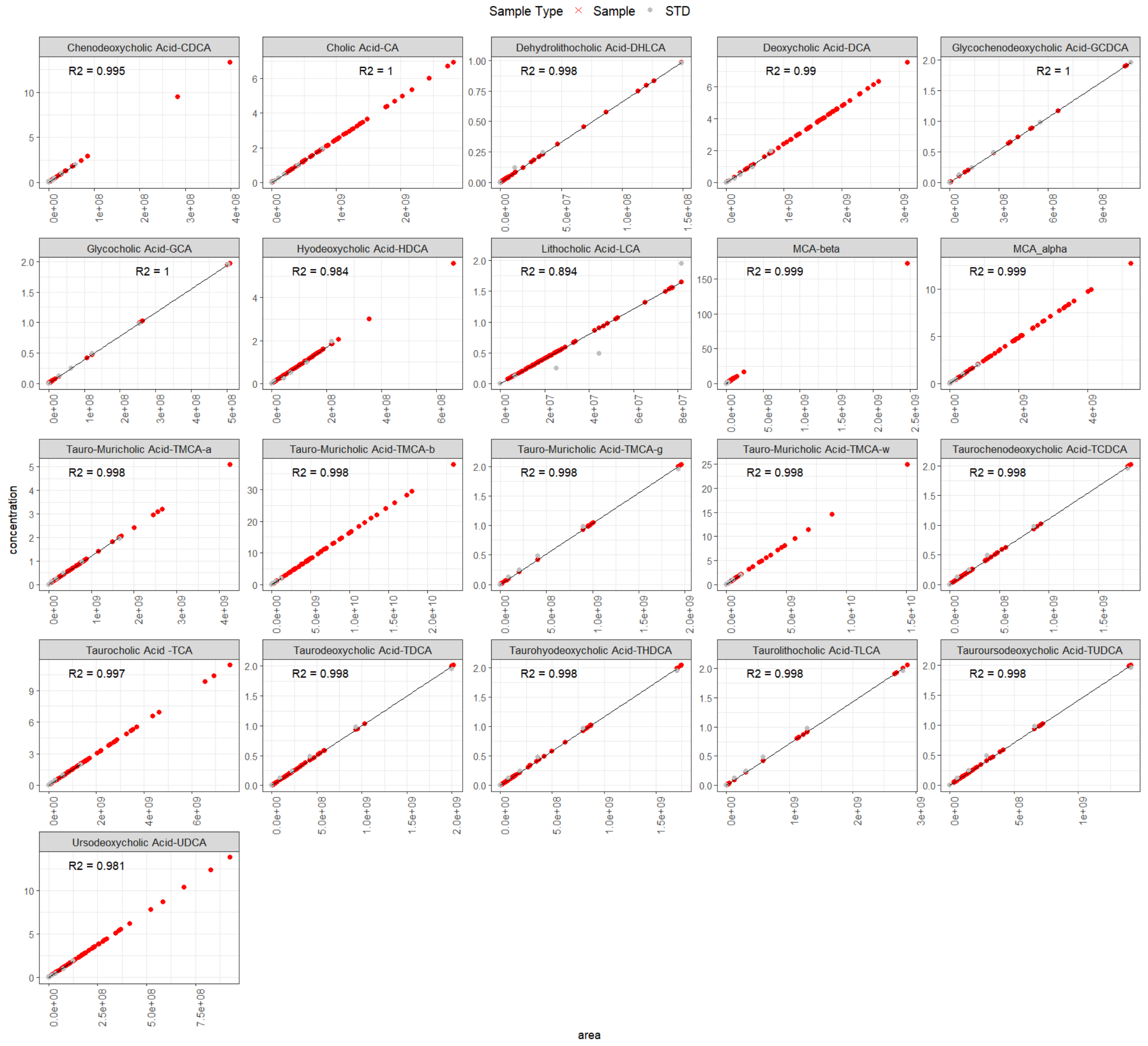


**Supplementary Figure 1: Standard curves for Fecal bile acid standards used for targeted quantification.**

Calibration curves generated from authentic bile acid standards demonstrating linearity across the dynamic range used for fecal bile acid quantification.

A

| RT (min) | CCS | Compound Name | Formula | Validated |
|---|---|---|---|---|
| 21.9 | 205.38743 | Dehydrolithocholic Acid-DHLCA | C24H38O3 | No |
| 21.9 | 202.820324 | Lithocholic Acid-LCA | C24H40O3 | No |
| 20.3 | 201.566797 | Hyodeoxycholic Acid-HDCA | C24H40O4 | Yes |
| 20.3 | 199.858695 | Ursodeoxycholic Acid-UDCA | C24H40O4 | Yes |
| 21.1 | 203.274907 | Chenodeoxycholic Acid-CDCA | C24H40O4 | Yes |
| 21.2 | 202.235188 | Deoxycholic Acid-DCA | C24H40O4 | No |
| 18.4 | 203.898137 | Muricholic Acid-MCA-alpha | C24H40O5 | Yes |
| 19.1 | 202.488939 | Muricholic Acid-MCA-beta | C24H40O5 | Yes |
| 20.2 | 202.859795 | Cholic Acid-CA | C24H40O5 | Yes |
| 18.1 | 201.078505 | Glycohyodeoxycholic Acid-GHDCA | C26H43NO5 | Yes |
| 18.1 | 200.930601 | Glycoursodeoxycholic Acid-GUDCA | C26H43NO5 | No |
| 20.3 | 200.486907 | Glycochenodeoxycholic Acid-GCDCA | C26H43NO5 | No |
| 18.1 | 201.908997 | Glycocholic Acid-GCA | C26H43NO6 | Yes |
| 21.4 | 206.343789 | Taurolithocholic Acid-TLCA | C26H45NO5S | Yes |
| 14 | 207.489119 | Tauroursodeoxycholic Acid-TUDCA | C26H45NO6S | Yes |
| 14.3 | 207.857782 | Taurohyodeoxycholic Acid-THDCA | C26H45NO6S | Yes |
| 18.9 | 206.973 | Taurochenodeoxycholic Acid-TCDCA | C26H45NO6S | Yes |
| 19.6 | 206.014496 | Taurodeoxycholic Acid-TDCA | C26H45NO6S | Yes |
| 6.3 | | Tauro-Muricholic Acid-TMCA-omega | C26H45NO7S | Yes |
| 6.8 | | Tauro-Muricholic Acid-TMCA-alpha | C26H45NO7S | Yes |
| 7.2 | | Tauro-Muricholic Acid-TMCA-beta | C26H45NO7S | Yes |
| 11.0 | | Tauro-Muricholic Acid-TMCA-gamma | C26H45NO7S | Yes |
| 14.9 | 207.317349 | Taurocholic Acid -TCA | C26H45NO7S | Yes |

B

| | Narrow (lower limit) | Wide (Upper limit) |
|---|---|---|
| m/z (ppm) | 2.0 | 5.0 |
| RT (min) | 0.1 | 0.5 |
| MSigma | 50 | 1000 |
| MS/MS score | 800 | 400 |
| CCS (%) | 1.0 | 5.0 |

**Supplementary Table S7. Liver bile acid standards and alignment criteria. (A)** Comprehensive list of bile acid standards provided by the Emory Integrated Metabolomics and Lipidomics Core (EIMLC) used for targeted liver bile acid quantification. **(B)** Alignment tolerances and scoring criteria applied for metabolite matching, including mass accuracy and retention time thresholds used to ensure confident feature annotation.

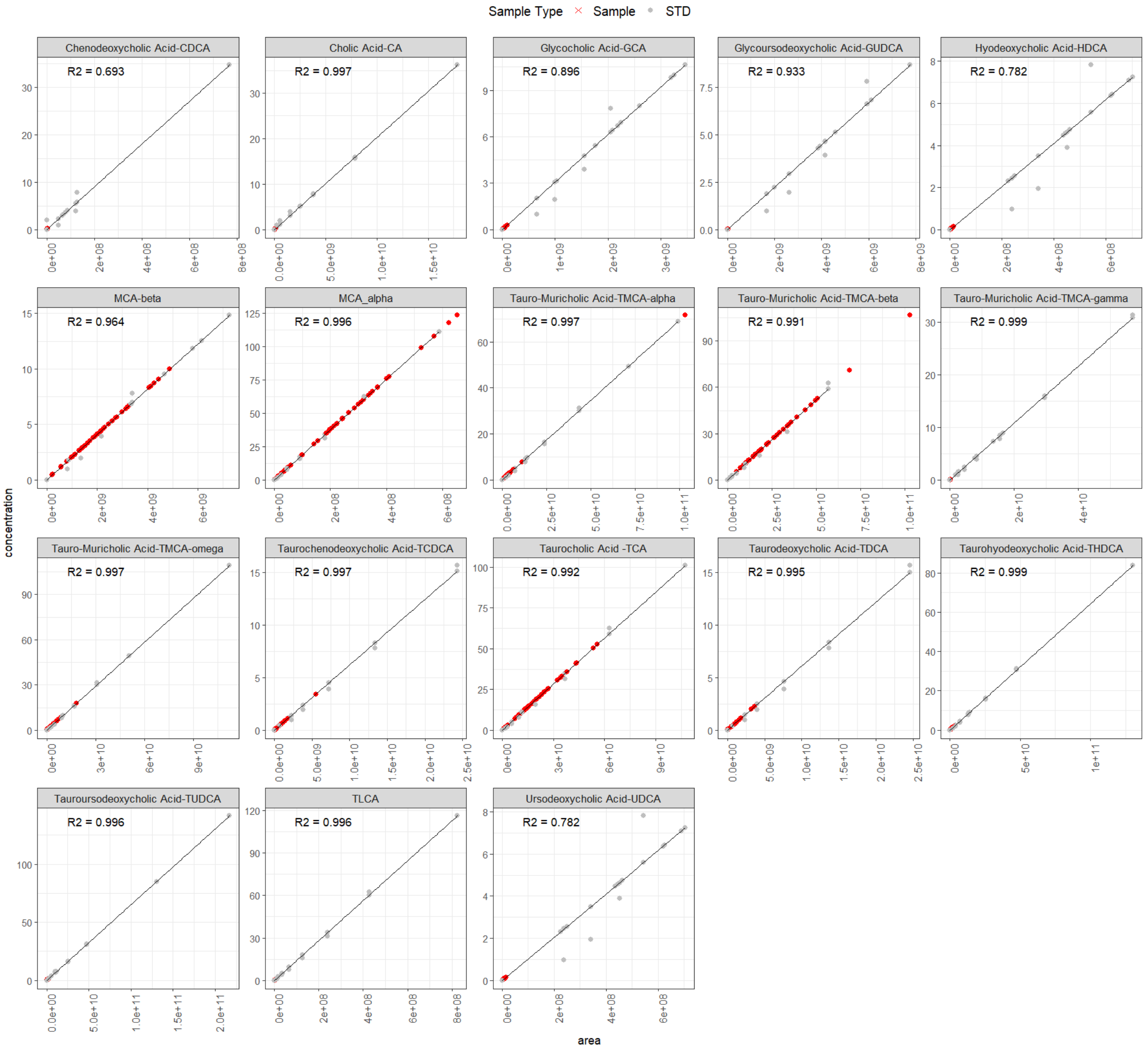


**Supplementary Figure 2: Standard curves for Liver bile acid standards used for targeted quantification.**

Calibration curves generated from authentic bile acid standards demonstrating linearity across the dynamic range used for liver bile acid quantification.

A

B

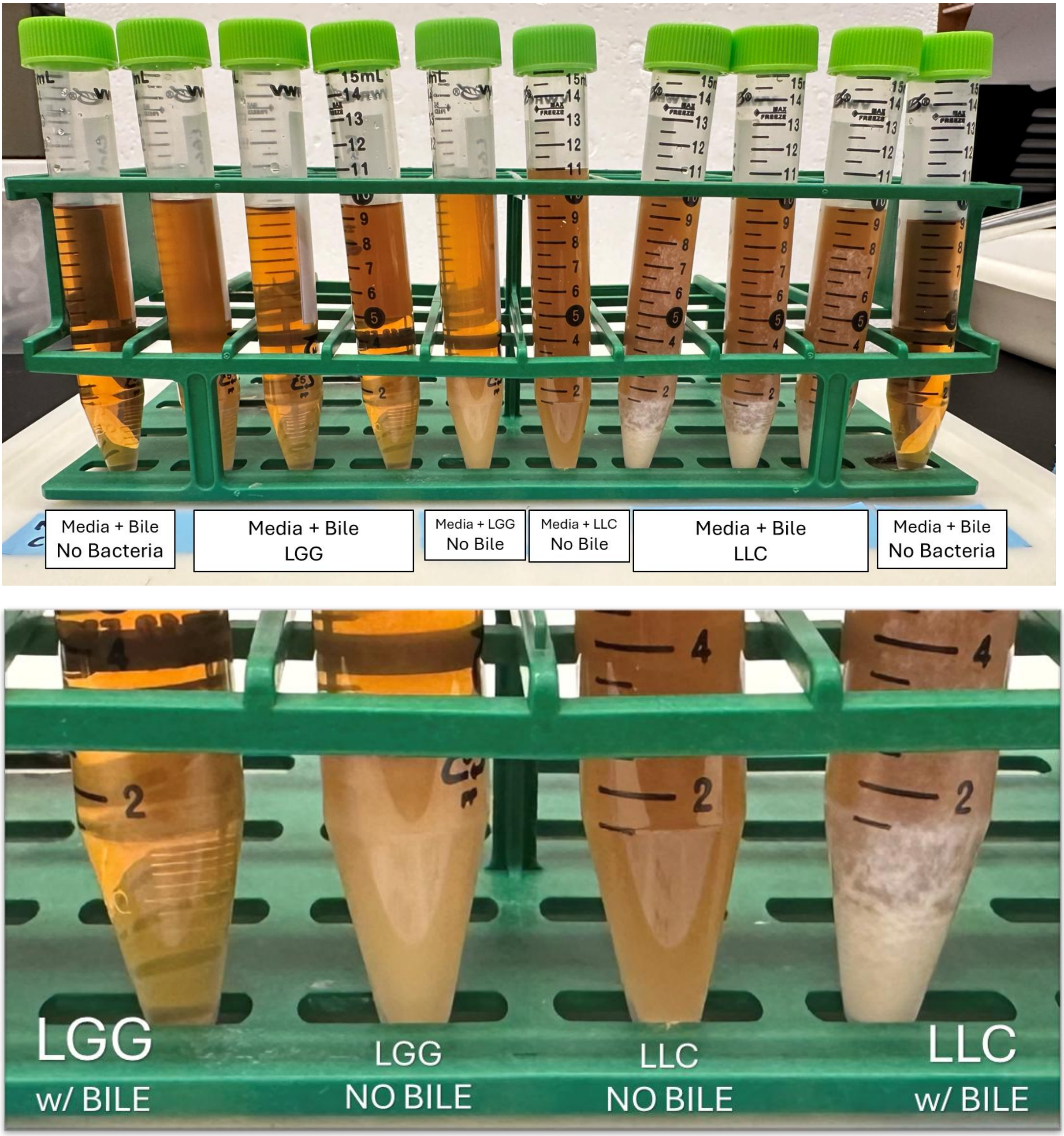


**Supplementary Figure 3. Qualitative bile salt hydrolase activity assessed using a pooled bile acid substrate.** **(A)** Representative image of all replicate culture conditions arranged from left to right as follows: MRS media supplemented with pooled bile acids but no bacteria (negative control); MRS + bile inoculated with *Lactobacillus rhamnosus* GG (LGG) (n = 3); MRS + LGG without bile (no–bile control); MRS + *Lactococcus lactis* subsp. cremoris (LLC) without bile (no–bile control); MRS + bile inoculated with LLC (n = 3); and an additional MRS + bile no-bacteria control. Visible precipitation in bile-supplemented cultures containing LGG or LLC is consistent with bile acid deconjugation, whereas no precipitation is observed in no–bile controls. **(B)** Representative close-up images of LGG and LLC cultures grown with or without pooled bile acids. Precipitation is evident in bile-supplemented cultures and is more pronounced in LLC compared to LGG under identical conditions, consistent with enhanced bile salt hydrolase activity.

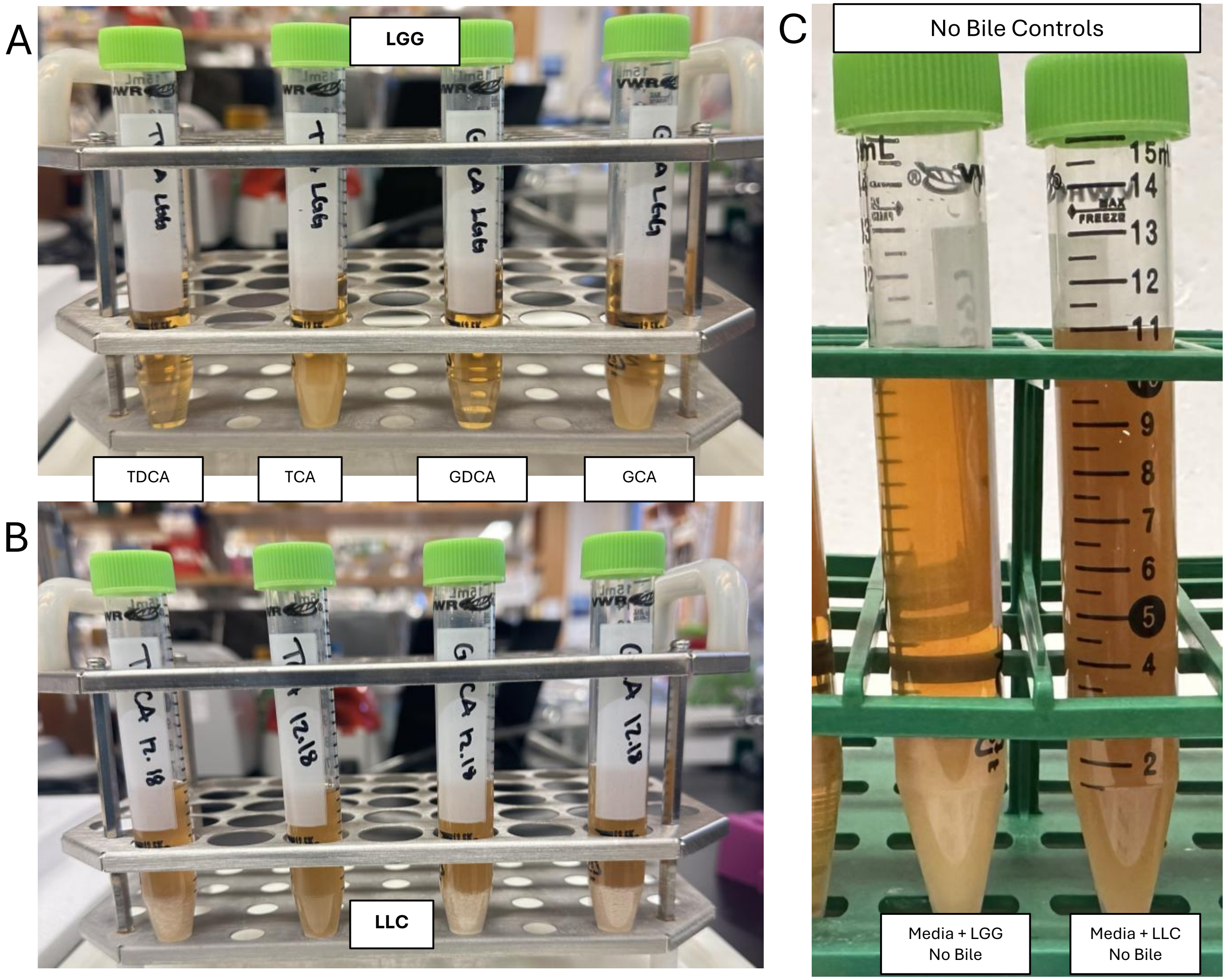


**Supplementary Figure 4. Qualitative bile salt hydrolase activity of LGG and LLC using individual bile acid substrates. (A)** *Lactobacillus rhamnosus* GG (LGG) cultures grown in MRS media supplemented individually with taurodeoxycholic acid (TDCA), taurocholic acid (TCA), glycodeoxycholic acid (GDCA), or glycocholic acid (GCA) (left to right; representative images shown). Visible precipitation in bile-supplemented cultures is consistent with bile acid deconjugation. **(B)** *Lactococcus lactis* subsp. cremoris (LLC) cultures grown under identical conditions with individual bile acids (TDCA, TCA, GDCA, GCA; left to right; representative images shown). Precipitation is observed across substrates and appears more pronounced in LLC cultures relative to LGG. **(C)** Representative close-up images of no–bile controls for LGG and LLC cultured in MRS media without bile acids, demonstrating absence of precipitation.

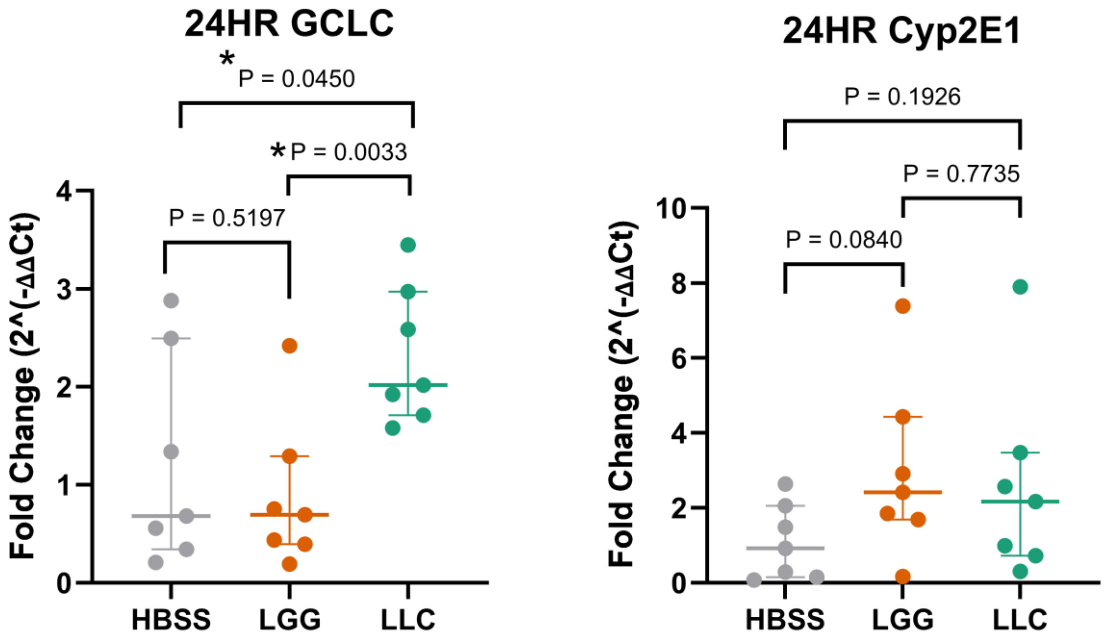


**Supplementary Figure 5. Regulation of oxidative stress and xenobiotic metabolism genes in HepG2 cells following exposure to bacterial culture supernatant. (A)** Relative mRNA expression of *GCLC* in HepG2 cells following 24-hour exposure to filter-sterilized bacterial culture supernatant. **(B)** Relative mRNA expression of *CYP2E1* under identical treatment conditions. Gene expression was quantified by RT–qPCR and normalized to housekeeping control 18s.
